# Biochemical selection on conserved glycans yield a cross-species AAV capsid for efficient panretinal intravitreal delivery

**DOI:** 10.64898/2026.11.17.767272

**Authors:** Jiang-Hui Wang, Mengtian Cui, Wenting Zhao, Delaney Giguere, Yue Yang, Qin Su, Ran He, Jun Xie, Qing Xie, Xiao Ke, Johanna M. Seddon, Claudio Punzo, Guangping Gao

## Abstract

Inherited retinal diseases are leading candidates for gene therapy, but photoreceptor-directed adeno-associated virus (AAV) vectors still require subretinal injection, which treats only part of the retina and carries surgical risk. Intravitreal delivery is simple and panretinal, yet inner-retinal barriers block most capsids, and capsids selected in one species rarely translate. Mapping retinal glycans across seven mammals, we found heparan sulfate (HS) marking the inner limiting membrane and terminal N-acetylglucosamine (GlcNAc) enriched on photoreceptor segments. These invariants defined a two-part design principle, attenuated HS binding with strong terminal GlcNAc engagement, implemented as a sequential in vitro selection, Conserved Receptor-Selection Technology (CReST). Its lead GlcNAc-selected capsid, AAV2.IvtB, increased retinal transgene expression 6.5-fold over AAV2.7m8 in mice and photoreceptor transduction 2.5-fold in pigs, acting through terminal GlcNAc engagement. It partially preserved photoreceptors in rd10 mice and suppressed laser-induced choroidal neovascularization. CReST is a receptor-guided, animal-free route to capsids with a defined, conserved mechanism.

**One-sentence summary:** A conserved-glycan selection run entirely in vitro yields an AAV capsid that reaches photoreceptors from the vitreous.

## Introduction

Inherited retinal diseases are among the most compelling targets for gene therapy: many arise from single-gene defects in the photoreceptors, the retina is immune-privileged and surgically accessible, and the approval of voretigene neparvovec for RPE65-associated disease established that AAV-based gene therapy can restore vision in humans.^1^ That success also exposed the field’s central constraint. Voretigene neparvovec is delivered by subretinal injection, in which vector is injected beneath the retina, raising a bleb that detaches the neurosensory retina from the underlying retinal pigment epithelium (RPE).^2^ The approach transduces photoreceptors and RPE efficiently, but reaches only the area covered by the bleb, requires invasive retinal surgery, and is poorly suited to the diffuse, panretinal degeneration typical of most inherited retinal disease.^3,4^

Intravitreal injection would resolve these limitations. It is a routine, low-risk office procedure, it distributes vector across the entire retina, and it is already the standard route for the anti-VEGF biologics used to treat millions of patients with neovascular retinal disease.^2^ The obstacle is biological: a vector placed in the vitreous must cross the inner retina to reach the photoreceptors of the outer retina, and the inner limiting membrane (ILM) together with the inner-retinal matrix forms a barrier that sequesters most AAV capsids before they arrive.^5–7^ Naturally occurring AAV2, the most-used retinal serotype, is detained at the inner retina by its affinity for heparan sulfate and rarely reaches photoreceptors after intravitreal delivery.^8,9^

Engineered capsids such as AAV2.7m8 have improved intravitreal transduction of the inner retina and, more modestly, of photoreceptors, and have advanced to clinical testing.^10^ Two limitations have nonetheless proven difficult to overcome. First, gains in the small, accessible mouse eye frequently shrink in larger eyes, where greater diffusion distances and a more substantial ILM impose stiffer barriers, an attrition that has repeatedly undermined translation.^10,11^ Second, most capsids are discovered by selecting for transduction directly in animals, a process that is slow and costly and that optimizes performance against the idiosyncrasies of a single host rather than against features shared with humans.^12,13^ Whether an engineered tropism translates depends on whether the molecular targets it exploits are themselves conserved, a question that outcome-based animal selection does not address.

We reasoned that a capsid able to reach photoreceptors across species should be engineered against the conserved molecular features of the intravitreal route rather than selected for an outcome in one host. Cell-surface glycans are the natural candidates for such targets, for three reasons. First, they are the primary attachment determinants for essentially every AAV serotype characterized to date, heparan sulfate for AAV2, sialic acid for AAV1, AAV5 and AAV6, galactose for AAV9,^14–18^ and attachment, rather than downstream trafficking, is the step at which the intravitreal route fails.^9,19,20^ Second, glycans encode spatial information that the protein components of AAV entry lack. The universal entry cofactors AAVR and GPR108 act intracellularly and are not known to be laminarly restricted within the retina;^21–23^ they therefore discriminate neither the barrier a vector must traverse nor the target cell it must reach. Glycan epitopes, by contrast, are laminarly patterned across the retina. Third, the two glycans that define this route are generated by core biosynthetic pathways shared across mammals, heparan sulfate as a ubiquitous basement-membrane glycosaminoglycan and N-acetylglucosamine (GlcNAc) as the invariant monosaccharide of every N-linked glycan, presented as an exposed terminal residue wherever those chains are incompletely elongated, so their retinal distribution can be measured directly in each species of interest rather than inferred from the sequence and expression differences that separate protein entry factors between rodents, pigs and primates, precisely the divergence most likely to erode an engineered tropism as it moves from the mouse eye to large-eye species. Glycan binding is also the property most tractable to capsid engineering,^24,25^ being governed by defined surface residues and measurable without animals using soluble competing sugars and glycosylation-deficient cell lines.

We therefore mapped the retinal distribution of the relevant cell-surface glycans across seven mammalian species and found a conserved pattern: heparan sulfate concentrated at the ILM, the barrier a vector must escape, and terminal GlcNAc enriched on the photoreceptor segments, the destination it must reach. These distributions define a two-part design principle: weakened heparan-sulfate binding to avoid sequestration, and enhanced terminal GlcNAc engagement to drive uptake at the segments, both anchored to conserved targets. Attenuation rather than ablation is the operative requirement on the heparan-sulfate side: the same interaction that sequesters capsid at the ILM is what first brings an intravitreally delivered vector to the vitreoretinal junction, and AAV2 variants in which heparan-sulfate binding is abolished fail to transduce the retina from the vitreous at all. The objective is therefore to rebalance the two interactions, not to remove one. Guided by this principle, we developed a receptor-guided, entirely *in vitro* selection platform and used it to derive AAV2.IvtB, a capsid that enhances photoreceptor transduction after intravitreal delivery in both mouse and pig, acts through a defined terminal GlcNAc-dependent mechanism, shows lower anti-AAV2 neutralizing-antibody titres than its parent in pigs, and improves structural and vascular outcomes in models of inherited photoreceptor degeneration and choroidal neovascularization.

## Results

### Conserved retinal localization of HS and terminal GlcNAc defines a targeting principle for intravitreal delivery

A capsid intended to work across species should engage molecular targets that are themselves conserved. We focused on cell-surface glycans because they, rather than protein receptors, govern the initial attachment step of nearly all AAV serotypes, because the glycosyltransferase pathways that generate them are conserved across mammals, and because their laminar distribution within the retina can specify both the barrier a vector must escape and the cell it must reach. Rather than screening for transduction in a single host, we therefore began by characterizing the surface glycans that an intravitreally delivered vector must encounter on its route from the vitreous to the photoreceptors. This route presents three sequential barriers (Fig. 1A): retention within the vitreous, the HS-rich ILM and inner-retinal matrix, and the receptor environment that governs cell entry across the inner and outer retina. Naturally occurring AAV2 is retained at the inner retina through HS binding and rarely reaches the outer layers.^9,14,24^ These observations defined the two capsid properties we set out to engineer.

**Fig 1.**
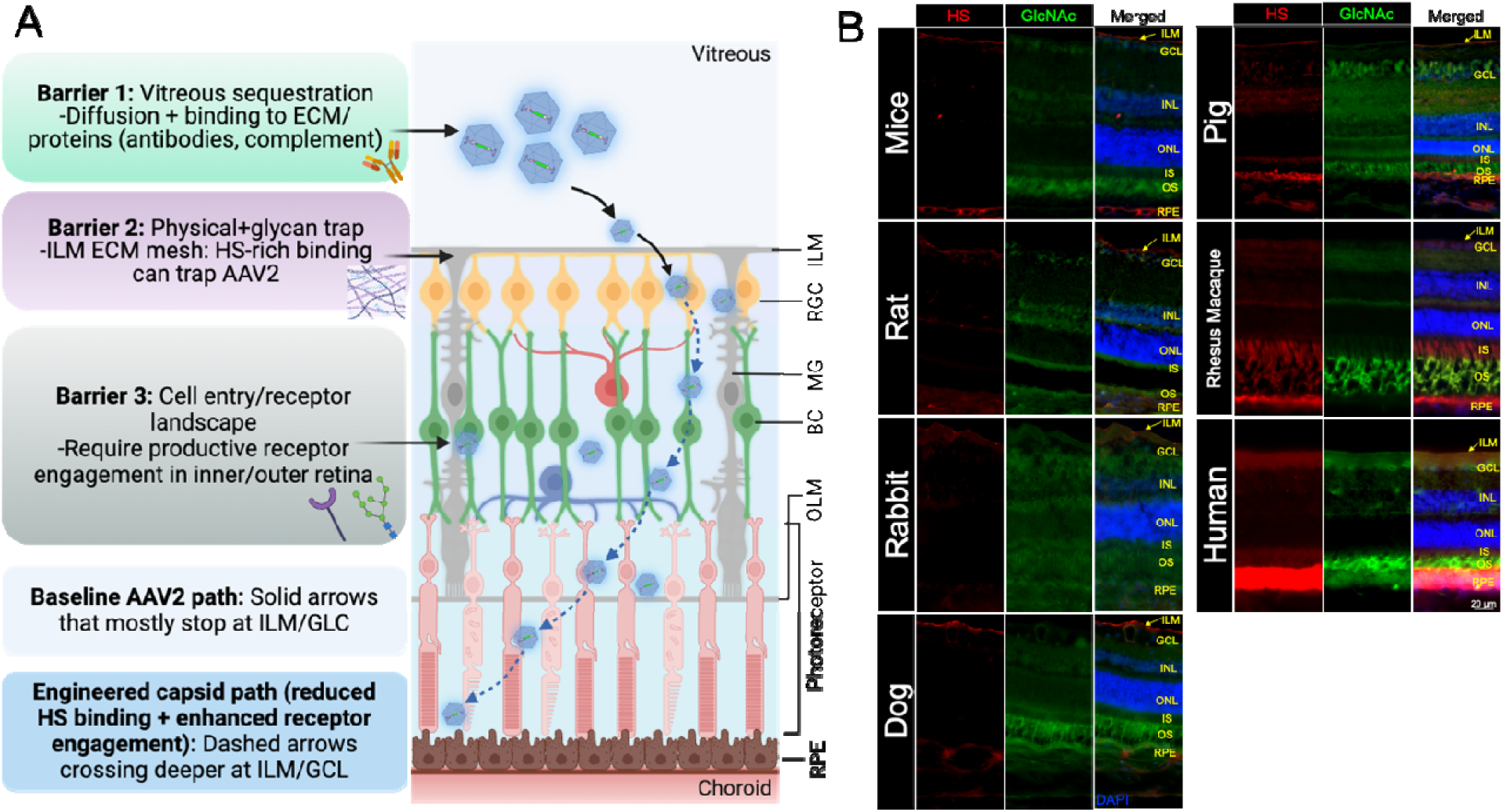
A conserved retinal glycan landscape defines receptor principles for intravitreal photoreceptor targeting. (A) Schematic of the barriers to intravitreal photoreceptor transduction (vitreous sequestration; the HS-rich ILM/extracellular-matrix trap; and the cell-entry/receptor landscape of the inner and outer retina) and the proposed engineered-capsid path (reduced HS binding plus enhanced receptor engagement) relative to the baseline AAV2 path. (B) Immunohistochemical localization of heparan sulfate (HS, red) and terminal N-acetylglucosamine (GlcNAc, green) with DAPI (blue) in retinal cross-sections from mouse, rat, rabbit, dog, pig, rhesus macaque, and human. HS is enriched at the ILM with weak inner-retinal and photoreceptor signal, whereas terminal GlcNAc is abundant on photoreceptor segments and conserved across species. Scale bar, 20 μm. GCL, ganglion-cell layer; ILM, inner limiting membrane; INL, inner nuclear layer; ONL, outer nuclear layer; IS/OS, inner/outer segments; RPE, retinal pigment epithelium.

We next examined whether the relevant glycan distributions are conserved. Using immunohistochemistry, we compared the retinal localization of HS and GlcNAc in seven mammalian species ranging from mouse to human (Fig. 1B). HS labeling was concentrated at the ILM and the RPE, with weak signal on inner-retinal neurons and photoreceptors. GlcNAc labeling was broader, with diffuse signal in the ganglion cell and inner nuclear layers, but its strongest and most discrete signal localized to the photoreceptor inner and outer segments, with little labeling in the outer nuclear layer. This segmental enrichment was most pronounced in the primate retinas (rhesus macaque and human). Two features relevant to intravitreal delivery were consistent across all seven species: an HS-rich ILM, and photoreceptor segments that carried the strongest GlcNAc labeling.

The barrier anatomy and the glycan distributions together define a two-part requirement for intravitreal photoreceptor targeting. An effective capsid should bind HS weakly, to avoid sequestration at the ILM, and engage a conserved, photoreceptor-enriched moiety such a terminal GlcNAc with sufficient avidity to drive productive uptake at the segments. Because both properties are anchored to glycans with conserved retinal localization, a capsid engineered around them should retain activity across species more reliably than one selected for function in a single host.

### Conserved Receptor-Selection Technology (CReST) yields AAV2.IvtB without animal screening

Guided by this principle, we designed CReST as a two-stage, receptor-guided selection carried out entirely *in vitro* (Fig. 2A). The input was an AAV2 peptide-display library carrying a random 12-mer insertion in the surface-exposed VR-VIII loop (after residue N587). The first stage selected against HS dependence. We placed an HS-coated Transwell membrane over a HeLa monolayer so that capsids with strong HS binding were preferentially retained by the barrier while those with weaker HS binding passed more readily to transduce the cells below. The membrane coating was optimized so that the system discriminates strongly from weakly HS-binding capsids (Fig. S1); transducing variants were recovered by reverse transcription of capsid-derived mRNA, depleting strongly HS-binding capsids from the pool. In the second stage, the HS-reduced library was panned on immobilized GlcNAc, and bound variants were recovered by genome extraction and quantified by next-generation sequencing (NGS). Lead candidates were then advanced to intravitreal validation in mice and pigs.

**Fig. 2.**
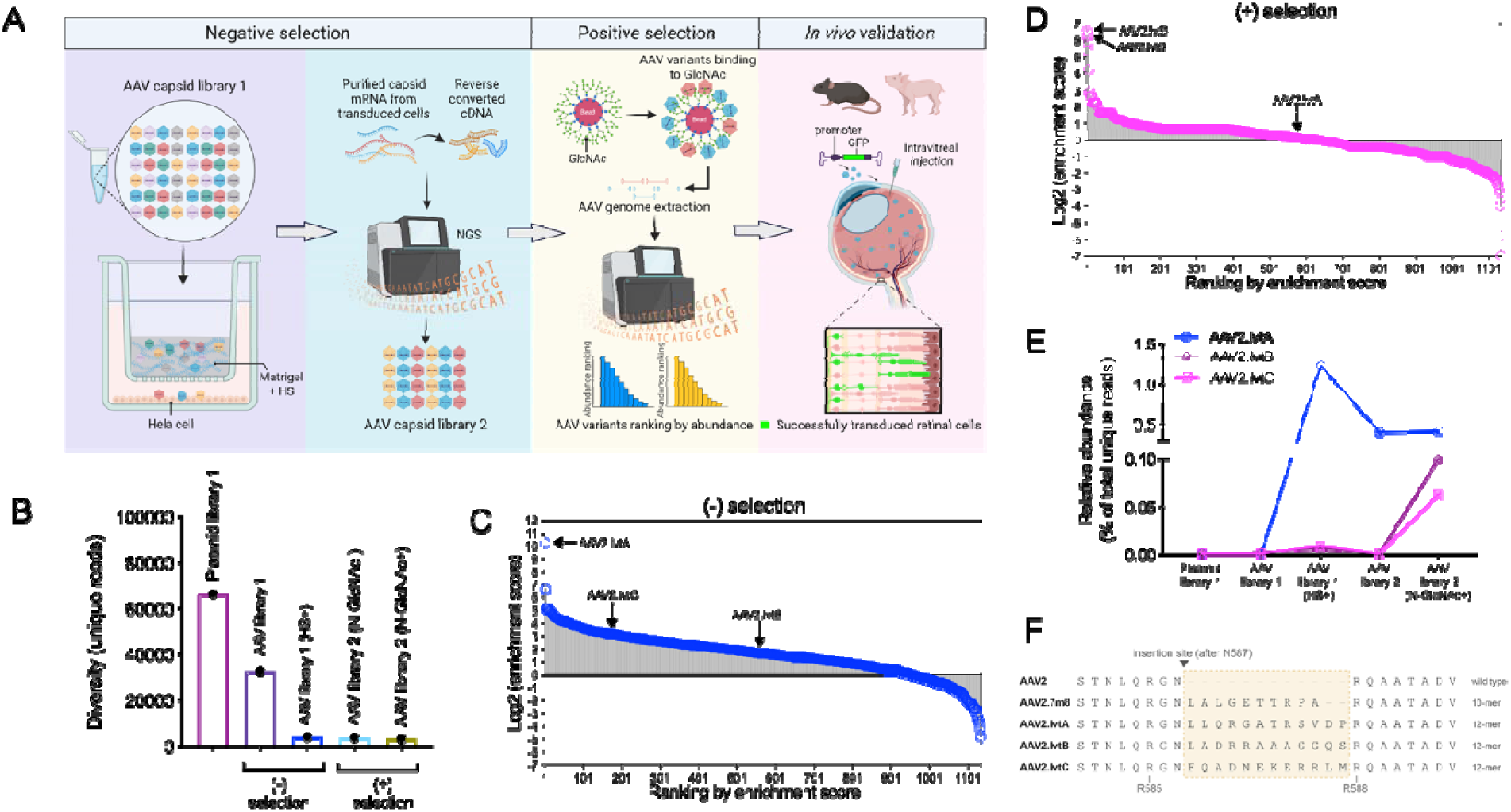
Conserved Receptor-Selection Technology (CReST) yields AAV2.IvtB without animal screening. (A) Workflow: negative selection on an HS-presenting substrate to de-enrich HS-dominated variants (capsid mRNA recovered from transduced cells), positive selection for GlcNAc binding, and intravitreal validation in mice and pigs. (B) Library diversity (unique reads) across selection stages, showing progressive collapse from the plasmid library through the HS-de-selected and GlcNAc-enriched pools. (C) Variant ranking by enrichment score in the HS-associated arm; AAV2.IvtA ranks highly. (D) Variant ranking in the GlcNAc arm; AAV2.IvtB and AAV2.IvtC are enriched. (E) Relative abundance of AAV2.IvtA, AAV2.IvtB and AAV2.IvtC across selection stages. Enrichment score: calculated as the relative RNA abundance of each variant in the transduced cells (Fig. 2C) or as the relative genomic DNA abundance of each variant after GlcNAc selection (Fig. 2D) normalized to its genomic abundance in the viral stock. (F) Amino acid sequence alignment at the VR-VIII insertion site (after residue N587) comparing wild-type AAV2 and AAV2.7m8 control, and the 12-mer inserts of AAV2.IvtA, AAV2.IvtB and AAV2.IvtC.

Library diversity contracted progressively through the selection, from 58,176 distinct 12-mer peptides in the plasmid library to 3,240 in the HS-reduced pool and 2,463 in the GlcNAc-enriched pool at comparable sequencing depth (91,360, 79,956 and 62,295 filtered reads; Fig. 2B). Sampling coverage rose across the same stages, indicating a genuine loss of richness rather than uneven depth. When variants were ranked by enrichment score, the two stages favored different capsids (Fig. 2C,D): AAV2.IvtA scored highly in the HS-based selection (Table S1), whereas AAV2.IvtB and AAV2.IvtC were most enriched in the GlcNAc-based selection (Table S2). Tracking individual variants across the two stages resolved three trajectories (Fig. 2E). AAV2.IvtA peaked in the HS-selected stage, AAV library 1 (HS□), and then declined to a lower, stable level, with no further gain in the GlcNAc pool. AAV2.IvtB remained near baseline through the HS stages and rose sharply only in the final GlcNAc-selected stage, AAV library 2 (GlcNAc□), where it became the most abundant variant. AAV2.IvtC followed the same trajectory as AAV2.IvtB, present at low abundance through the HS stages and rising only in th GlcNAc-selected pool, but reached a lower final level. Alignment across the VR-VIII insertion site confirmed that AAV2.IvtA, AAV2.IvtB and AAV2.IvtC each carry a different 12-mer peptide at this position, distinct from wild-type AAV2 and from the 10-mer AAV2.7m8 control (Fig. 2F). We carried AAV2.IvtB and AAV2.IvtC forward as the lead GlcNAc-selected capsids and retained AAV2.IvtA as a comparator from the HS-negative selection.

### AAV2.IvtB enhances pan-retinal and photoreceptor transduction after intravitreal delivery in mice

We intravitreally delivered single-stranded rAAV expressing GFP from a CMV-enhancer/chicken β-actin (CB6) promoter at 2×10^9^ vector genomes (vg) per eye, and compared the engineered capsids with parental AAV2 and AAV2.7m8. Four weeks after injection, fundus fluorescence imaging showed broad, bright retinal GFP for AAV2.IvtA, AAV2.IvtB and AAV2.IvtC, whereas AAV2 gave only sparse signal and AAV2.7m8 a predominantly perivascular signal (Fig. 3A). Retinal GFP transcript levels were highest for AAV2.IvtB (≈6.5-fold over AAV2.7m8; P < 0.0001) and AAV2.IvtA (P = 0.0009); AAV2.IvtC was intermediate and not significantly above AAV2.7m8 (Fig. 3B). However, at molecular level, AAV2.IvtB carried the highest vector-genome load (P = 0.0012) at a transcript-per-genome ratio similar to AAV2.7m8, whereas AAV2.IvtA delivered fewer genomes but expressed more per genome (P = 0.0113).

**Fig. 3.**
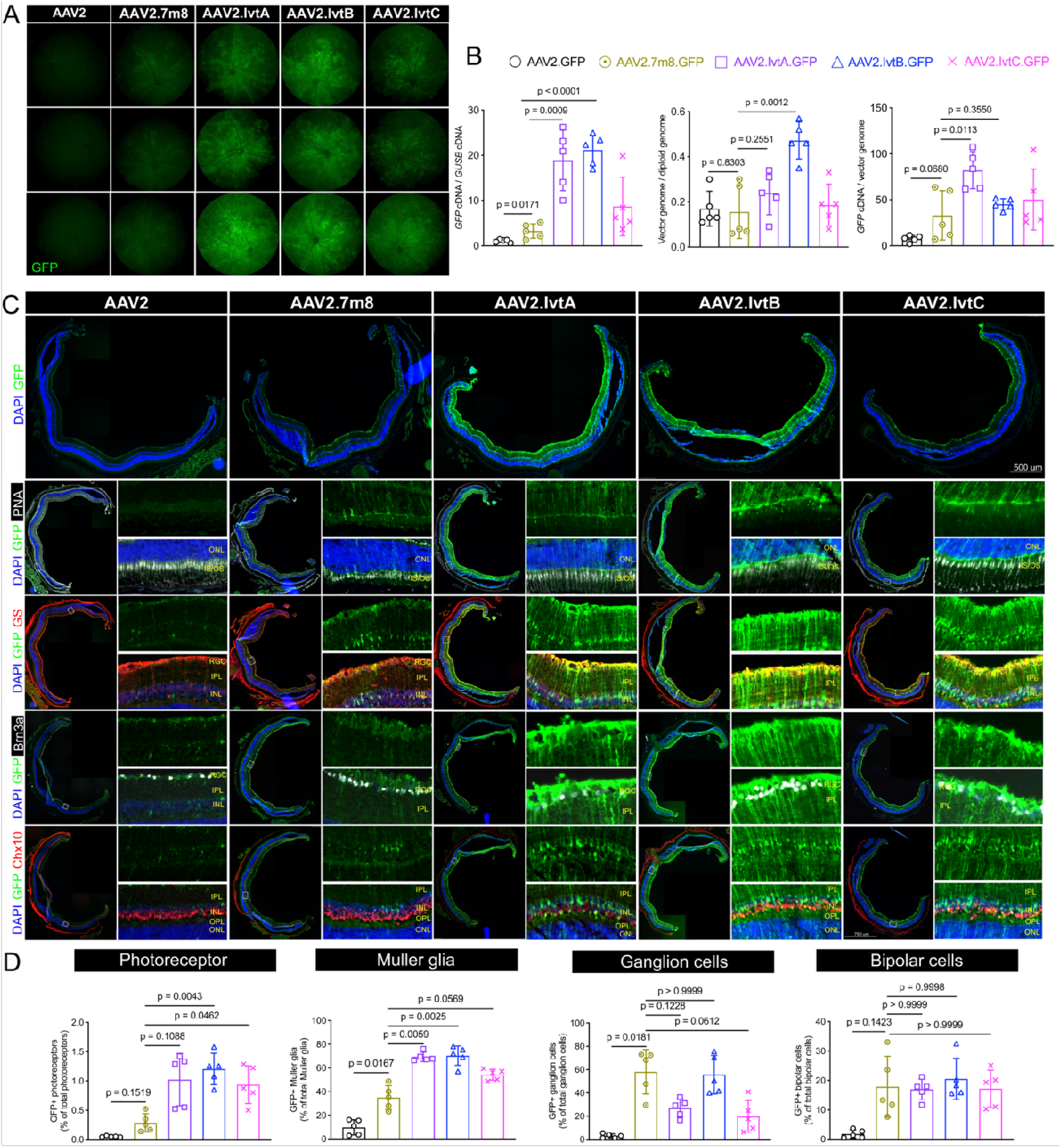
AAV2.IvtB enhances pan-retinal and photoreceptor transduction after intravitreal delivery in mice. Single-stranded CB6-GFP rAAV delivered intravitreally at 2×10^9^ vg/eye. (A) Representative fundus GFP fluorescence for AAV2, AAV2.7m8, AAV2.IvtA, AAV2.IvtB and AAV2.IvtC. (B) Retinal GFP transcripts (normalized to *Gusb*), vector genomes (per diploid genome) and GFP transcript per vector genome; AAV2.IvtB shows the highest expression (≈6.5-fold over AAV2.7m8 transcripts, p < 0.0001, n = 5 per group) and higher vector-genome abundance (p = 0.0012). (C) Retinal cross-sections co-stained for GFP with PNA (cone outer segments), GS (Müller glia), Brn3a (ganglion cells) and Chx10 (bipolar cells). (D) Cell-type quantification; AAV2.IvtB increases photoreceptor transduction 4.1-fold (p = 0.0043) and Müller glia 2.0-fold (p = 0.0025) versus AAV2.7m8 (n = 5 per group). Scale bar, 500 μm. Data are means ± SD. Each readout was compared across capsids by one-way ANOVA with Dunnett’s correction for multiple comparisons.

To determine cell-type tropism, we co-stained retinal cross-sections for markers of cone photoreceptors (peanut agglutinin, PNA), Müller glia (glutamine synthetase, GS), ganglion cells (Brn3a) and bipolar cells (Chx10) (Fig. 3C; Fig. S2). AAV2 transduced the retina only sparsely, and AAV2.7m8 was largely confined to the inner retina, with little signal in the photoreceptor segments. By contrast, AAV2.IvtA, AAV2.IvtB and AAV2.IvtC each transduced the full retinal thickness, showing the radial profile characteristic of Müller glia together with clear GFP signal in the photoreceptor segments; transduction was most extensive for AAV2.IvtB. Quantification confirmed these patterns (Fig. 3D). Photoreceptor transduction was scored as GFP-positive segments relative to the total outer nuclear layer nuclei in the same area; PNA labels cone outer segments and was used to visualize the segment layer rather than as the denominator. All three engineered capsids transduced photoreceptors more efficiently than AAV2.7m8, with the greatest effect for AAV2.IvtB (4.1-fold; P = 0.0043). AAV2.IvtB also gave the highest Müller glia transduction (2.0-fold over AAV2.7m8; P = 0.0025), and AAV2.IvtA and AAV2.IvtC were likewise elevated. Ganglion cell transduction was high for both AAV2.7m8 and AAV2.IvtB and lower for AAV2.IvtA and AAV2.IvtC, whereas bipolar cell transduction was comparable across transducing engineered capsids. AAV2.7m8 and AAV2.IvtB thus differed mainly in the outer retina, which AAV2.IvtB transduced far more effectively.

### The transduction advantage translates to the large-animal pig eye

Because the murine eye is small and anatomically distinct from the human eye, cross-species validation in a large eye is essential for translational interpretation.^5,6,10^ We therefore delivered the same GFP vectors intravitreally to pigs at 2×10^11^ vg per eye. *Ex vivo* imaging of dissected retina showed the brightest, most widespread retinal GFP for AAV2.IvtB, exceeding AAV2.7m8, AAV2.IvtA and parental AAV2 (Fig. 4A). Molecular quantification confirmed this: AAV2.IvtB produced the highest GFP transcript levels (p = 0.0114 versus AAV2; p = 0.0483 versus AAV2.7m8; Fig. 4B). Vector-genome abundance was comparable across all capsids (Fig. 4B). AAV2.IvtB showed the highest expression per genome (p = 0.0019 versus AAV2; p = 0.0335 versus AAV2.7m8; Fig. 4B).

**Fig. 4.**
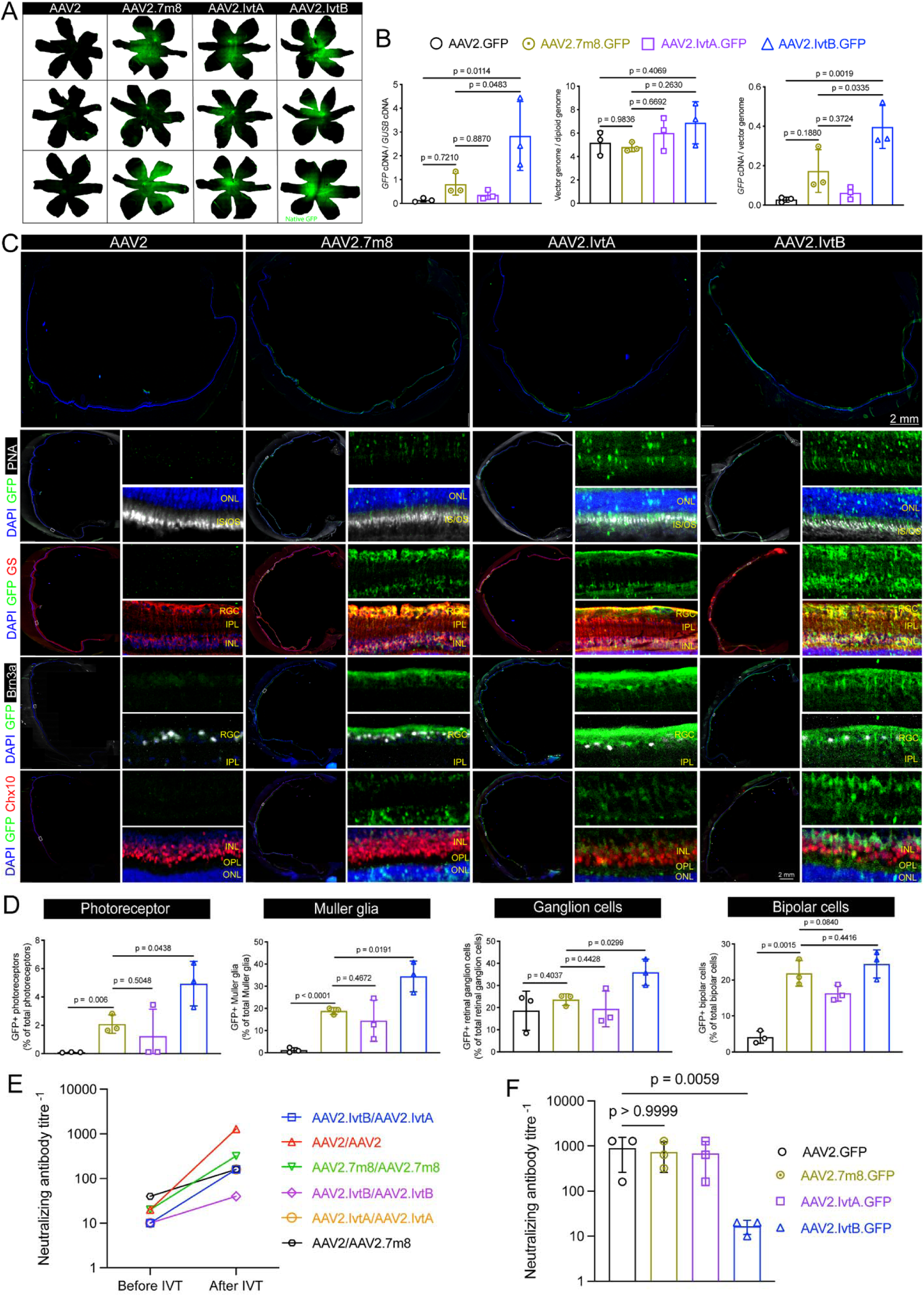
Enhanced AAV2.IvtB transduction translates to the pig eye with a lower anti-AAV2 neutralizing-antibody response. Single-stranded CB6-GFP rAAV delivered intravitreally to pigs at 2×10^11^ vg/eye. (A) Ex vivo imaging of dissected retina showing GFP fluorescence. (B) Retinal GFP transcripts, vector genomes and GFP per vector genome; AAV2.IvtB is highest (p = 0.0114 transcripts; p = 0.0019 transcripts per genome versus AAV2, n = 3 per group). (C) Cross-sections co-stained for GFP with PNA (cone outer segments), GS (Müller glia), Brn3a (ganglion cells) and Chx10 (bipolar cells). Quantification was performed as described in Materials and Methods. (D) Cell-type quantification; AAV2.IvtB increases photoreceptor 2.5-fold (p = 0.0438), Müller glia 1.7-fold (p = 0.0191) and ganglion-cell 1.5-fold (p = 0.0299) transduction versus AAV2.7m8, with no difference in bipolar cells (p = 0.4416) (n = 3 per group). (E) Serum anti-AAV2 NAb titres before injection and at endpoint, plotted per animal and colored by the capsid injected in that animal’s eyes. Baseline titres were uniformly low (1:10–1:40). Among the four animals injected with the same capsid in both eyes, endpoint titres were highest for AAV2 (1:1,280), intermediate for AAV2.7m8 (1:320) and AAV2.IvtA (1:160), and lowest for AAV2.IvtB (1:40); two animals received different capsids in each eye and are shown for completeness but excluded from the per-capsid comparison. (F) Vitreous anti-AAV2 NAb titres at endpoint, grouped by injected capsid (n = 3 eyes per capsid): AAV2 and AAV2.7m8, 1:640; AAV2.IvtA, 1:508; AAV2.IvtB, 1:16 (≈40-fold lower than AAV2). Scale bar, 2 mm. Data are mean ± SD unless stated otherwise. Each readout was compared across capsids by one-way ANOVA with Dunnett’s correction for multiple comparisons.

Cross-sections co-stained for photoreceptors, Müller glia, ganglion cells and bipolar cells showed that AAV2.IvtB transduced the full retinal thickness, with the clearest photoreceptor-segment (IS/OS) and Müller signal (Fig. 4C; Fig. S3). Quantification (Fig. 4D) showed that AAV2.IvtB significantly increased photoreceptor (2.5-fold; p = 0.0438), Müller-glia (1.7-fold; p = 0.0191) and ganglion-cell (1.5-fold; p = 0.0299) transduction over AAV2.7m8, whereas bipolar-cell transduction did not differ between the two (p = 0.4416). The photoreceptor fold-improvement was smaller than in mice, as expected given the larger vitreal volume and longer diffusion distances of the porcine eye, but the direction and significance of the effect were preserved across species. Retinal transcript levels of inflammatory cytokines (IFN-γ, IL-1β, IL-6, TNF-α) did not differ significantly among capsids (Fig. S4), indicating that enhanced transduction was not accompanied by a detectable pro-inflammatory signature.

Beyond transduction, AAV2.IvtB also elicited a lower anti-AAV2 neutralizing-antibody (NAb) response than AAV2, AAV2.7m8 or AAV2.IvtA after intravitreal delivery in pigs (Fig. 4E). Before injection, anti-AAV2 NAb titres in serum were uniformly low across all animals (1:10–1:40). By endpoint, titres in the four animals injected with the same capsid in both eyes had risen substantially for AAV2 (1:1,280), were intermediate for AAV2.7m8 (1:320) and AAV2.IvtA (1:160), and remained lowest for AAV2.IvtB (1:40); because each of these serum values reflects a single animal, this comparison is descriptive rather than statistical. The vitreous, sampled independently from each eye (n = 3 eyes per capsid), showed the same ranking with better support: geometric-mean titres were 1:640 for both AAV2 and AAV2.7m8, 1:508 for AAV2.IvtA, and 1:16 for AAV2.IvtB, a ≈40-fold reduction relative to AAV2 (Fig. 4F).

### Intravitreal AAV2.IvtB improves structural and vascular outcomes in two distinct retinal disease models

Having established where AAV2.IvtB penetrates, how it enters cells and the humoral response it provokes, we asked whether these properties translate into therapeutic benefit in two models posing complementary requirements: gene supplement in inherited photoreceptor degeneration, and delivery of a secreted biologic in a vascular retinal disease. For the first, we used the rd10 mouse, a model of autosomal recessive retinitis pigmentosa in which a missense mutation in *Pde6b*, the gene encoding the β-subunit of rod cGMP phosphodiesterase, causes rapid rod photoreceptor degeneration, with the first histological changes at about postnatal day 16 and peak rod loss between P18 and P25, followed by secondary cone loss.^26,27^ We delivered *Pde6b* intravitreally at postnatal day 15, at the earliest point compatible with the injection procedure and immediately preceding the first detectable histological change, and compared AAV2.IvtB.*Pde6b* with AAV2.*Pde6b*, AAV2.7m8.*Pde6b*, an AAV2.GFP-treated control and untreated wild-type retina at P45 (Fig. 5A). In AAV2.GFP-treated rd10 retinas the outer nuclear layer (ONL) was severely thinned relative to wild-type, as expected for untreated degeneration at this age.^27^ *Pde6b* delivery slowed this loss, and the effect was greatest with AAV2.IvtB, whose ONL was visibly thicker than that of AAV2.*Pde6b* or AAV2.7m8.*Pde6b* (Fig. 5B). ONL thickness with AAV2.IvtB.*Pde6b* (19.6 µm) was greater than AAV2.*Pde6b* (10.3 µm, p = 0.0032) and AAV2.7m8.*Pde6b* (12.6 µm, p = 0.0085), but remained below the wild-type thickness (66 µm) measured in parallel (Fig. 5C). Because treatment began at postnatal day 15, when degeneration is already beginning and transgene expression has yet to reach steady state, and because structure was assessed at a single endpoint, these data do not distinguish slowed from arrested degeneration. What the comparison supports is that the greater ONL preservation achieved by AAV2.IvtB.*Pde6b* tracks with the higher photoreceptor transduction measured for AAV2.IvtB over AAV2.7m8 in wild-type retina. AAV2.IvtB.*Pde6b* also produced the highest retinal *Pde6b* transcript levels among treated groups, significantly above AAV2.*Pde6b* (p = 0.0263), without increased vector delivery (retinal vector-genome copies did not differ among treated capsids, p = 0.1727; Fig. 5D), consistent with the advantage deriving from more productive transduction rather than from greater vector delivery.

**Fig. 5.**
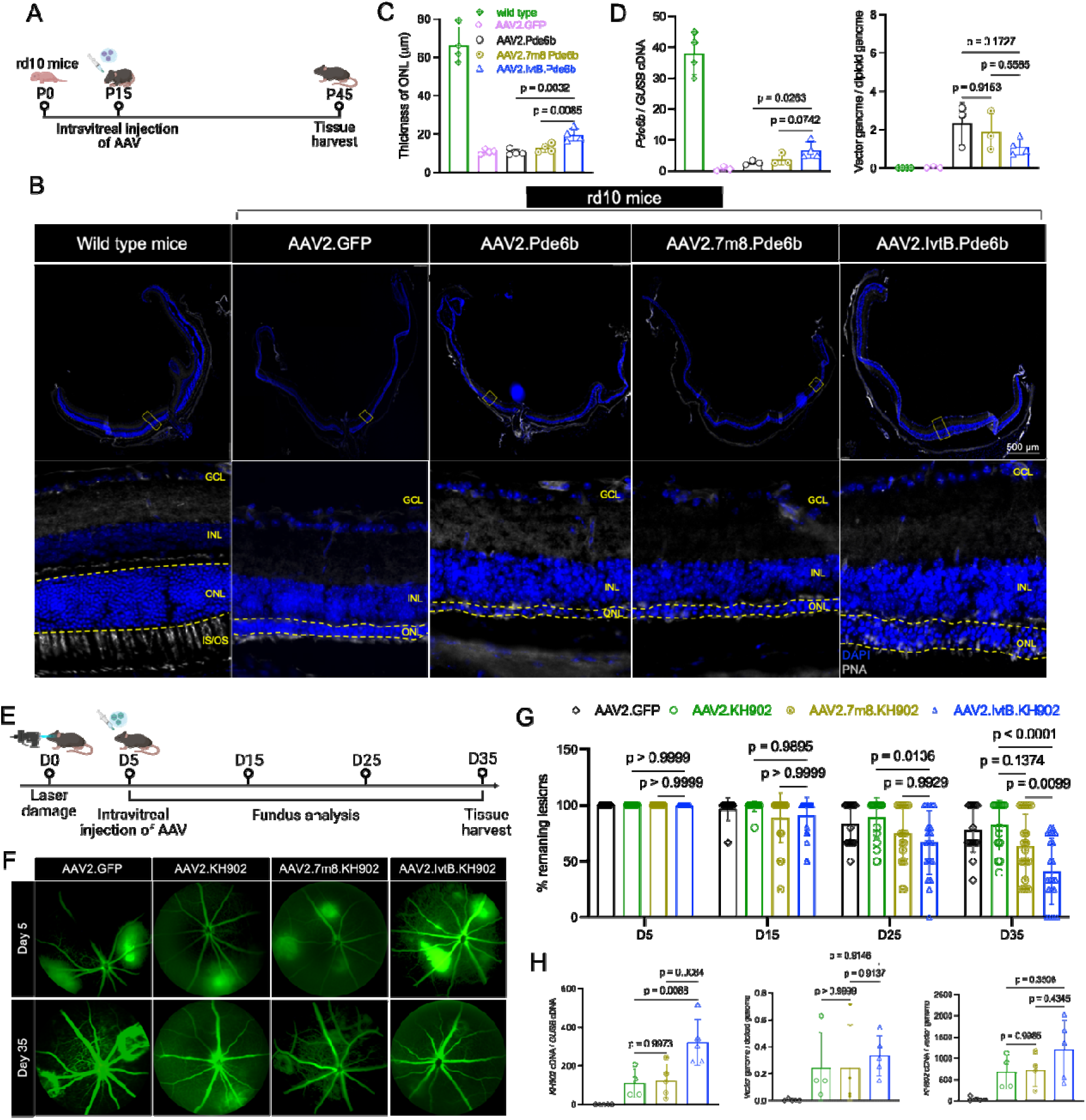
Intravitreal AAV2.IvtB improves structural and vascular outcomes in two distinct retinal disease models. (A) Design: rd10 mice received intravitreal AAV at postnatal day 15 with analysis at day 45. (B) Representative cross-sections (DAPI, PNA) for wild-type and rd10 retinas treated with AAV2.GFP, AAV2.*Pde6b*, AAV2.7m8.*Pde6b* or AAV2.IvtB.*Pde6b*. (C) ONL thickness; AAV2.IvtB.*Pde6b* partially preserves the ONL (p = 0.0032 versus AAV2.*Pde6b*; p = 0.0085 versu AAV2.7m8.*Pde6b;* n = 4 per group) but remains below wild-type. (D) Retinal *Pde6b* transcripts and vector genomes; AAV2.IvtB.*Pde6b* is higher than AAV2.*Pde6b* (p = 0.0263; n = 3-4 per group), with no difference in vector genomes among treated capsids. (E–H) Laser-induced choroidal neovascularization. (E) Design: laser photocoagulation on day 0, intravitreal AAV.KH902 on day 5, fundus fluorescein angiography through day 35. (F) Representative angiograms at day 5 and day 35. (G) Proportion of remaining CNV lesions over time; AAV2.IvtB.KH902 reduces lesions by day 35 versus AAV2.GFP (p < 0.0001) and AAV2.7m8.KH902 (p = 0.0099). (H) Retinal KH902 transcripts and vector genomes; AAV2.IvtB delivers ≈2.5-to 3-fold higher KH902 transcript (p = 0.0088 versus AAV2.KH902; p = 0.0084 versus AAV2.7m8.KH902) with no difference in vector genomes (n=4-5 per group). Scale bar, 500 μm. Data are mean ± SD unless stated otherwise. Each readout was compared across capsids by one-way ANOVA with Dunnett’s correction for multiple comparisons.

For the second model, we delivered the anti-VEGF agent KH902^28^ intravitreally in a laser-induced choroidal neovascularization (CNV) model and compared AAV2.IvtB.KH902 with AAV2.KH902, AAV2.7m8.KH902 and an AAV2.GFP control, tracking lesions by fundus fluorescein angiography from day 5 to day 35 (Fig. 5E). CNV was induced by laser photocoagulation on day 0 and vectors were delivered intravitreally on day 5, at which point the baseline lesion number that developed in each group was recorded. Fluorescein leakage from CNV lesions persisted in AAV2.GFP-treated eyes, whereas KH902 delivery progressively reduced lesion burden (Fig. 5F). Lesion burden did not differ among the groups at day 5, before vector was given, confirming balanced randomization, or at day 15, but AAV2.IvtB.KH902 diverged with time: by day 25 it had significantly fewer remaining lesions than AAV2.KH902 (p = 0.0136), and by day 35 the remaining lesion burden (≈40%) was lower than both the AAV2.GFP control (p < 0.0001) and AAV2.7m8.KH902 (p = 0.0099) (Fig. 5G). This benefit tracked with transgene expression: retinal KH902 transcript was ≈2.5-to 3-fold higher with AAV2.IvtB.KH902 than with AAV2.7m8.KH902 (p = 0.0084) or AAV2.KH902 (p = 0.0088), whereas retinal vector-genome abundance did not differ among the treated capsids (p ≥ 0.91; Fig. 5H). Together, these two models demonstrate benefit for both a gene-supplementation and a secreted-biologic modality, each driven by greater expression per genome rather than by more genomes.

### Enhanced terminal GlcNAc engagement drives AAV2.IvtB attachment and entry

The capsids had been selected for defined glycan-binding properties, but it remained to be shown which interaction actually drives cell entry. We therefore compared AAV2, AAV2.7m8 and AAV2.IvtB in parental CHO-K1 cells and in the isogenic mutant Lec8, which lacks the Golgi UDP-galactose transporter and therefore presents N-glycans truncated to terminal GlcNAc. Surface staining and flow cytometry confirmed that the two lines differ in terminal GlcNAc availability on a comparable heparan-sulfate background (Fig. S5). On the CHO-K1 background the three capsids were not separated by transduction: AAV2.IvtB matched parental AAV2 (p = 0.4551), and AAV2.7m8 transduced less well than either (p < 0.0001) (Fig. 6A,B). In Lec8, AAV2.IvtB transduced approximately twice as efficiently as AAV2 (p < 0.0001), whereas AAV2.7m8 remained low (p < 0.0001). The advantage of AAV2.IvtB is therefore conditional on an altered-glycan background rather than a general gain in infectivity.

**Fig. 6.**
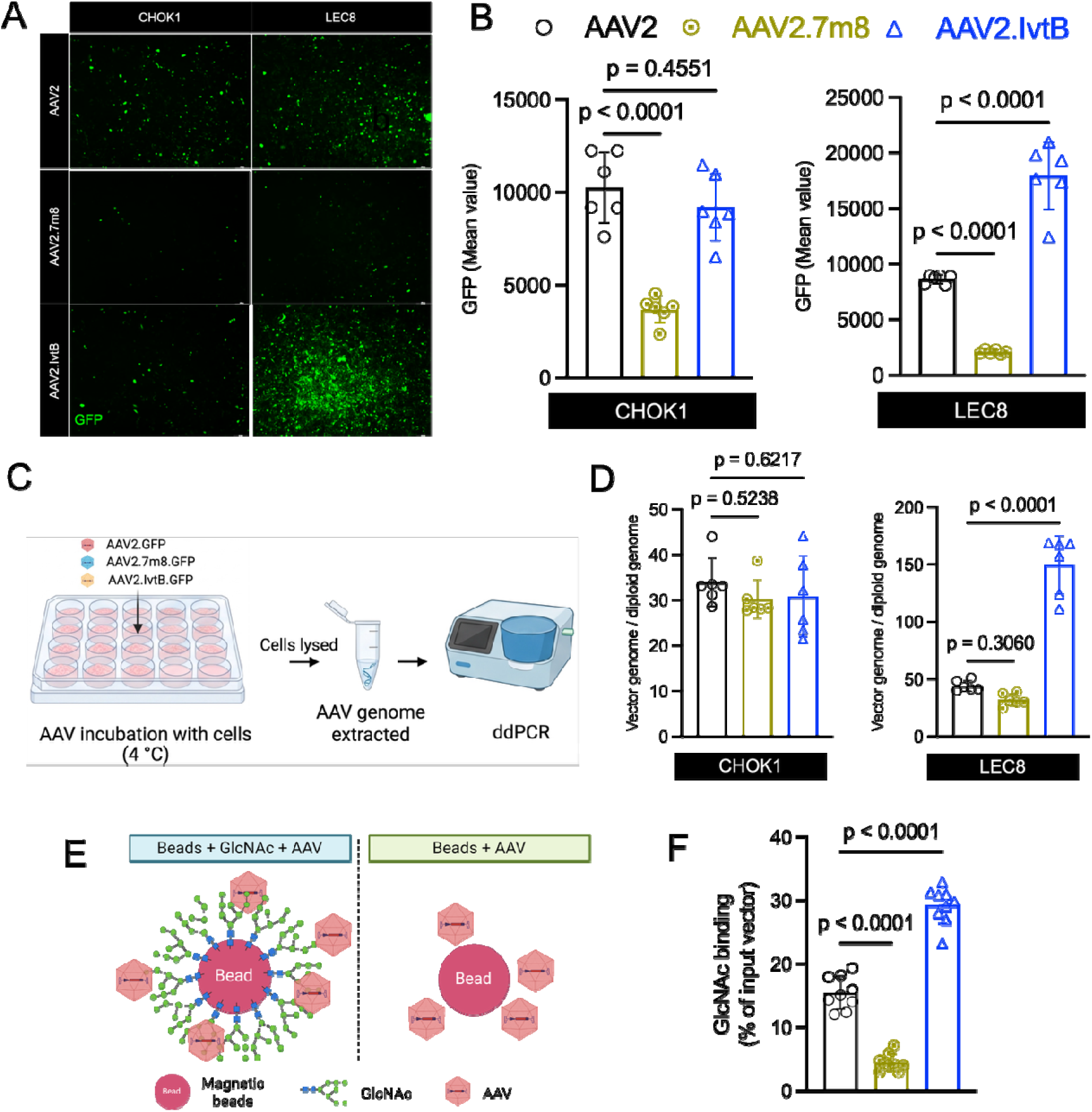
Enhanced terminal GlcNAc engagement drives AAV2.IvtB attachment and transduction. Throughout, CHO-K1 cells carry wild-type glycosylation and Lec8 cells lack the Golgi UDP-galactose transporter, presenting N-glycans truncated to terminal GlcNAc. (A) GFP fluorescence in CHO-K1 and Lec8 cells transduced with AAV2.GFP, AAV2.7m8.GFP or AAV2.IvtB.GFP. (B) Mean GFP fluorescence intensity measured by flow cytometry in parallel cultures. CHO-K1: AAV2.IvtB versu AAV2, p = 0.4551; AAV2.7m8 versus AAV2, p < 0.0001. Lec8: AAV2.IvtB versus AAV2, p < 0.0001; AAV2.7m8 versus AAV2, p < 0.0001; n = 6 per group. (C) Schematic of the cell-binding assay; vector was bound at 4 °C, cells were washed and lysed, and bound genomes were quantified by ddPCR. (D) Cell-bound vector genomes per diploid genome. CHO-K1: p = 0.6217 and p = 0.5238 versus AAV2. Lec8: AAV2.IvtB versus AAV2, p < 0.0001; AAV2.7m8 versus AAV2, p = 0.3060; n = 6 per group. (E) Schematic of the cell-free pull-down; magnetic beads with or without immobilized GlcNAc were incubated with vector. (F) Vector recovered on GlcNAc beads as a percentage of input. AAV2.IvtB versus AAV2, p < 0.0001; AAV2.7m8 versus AAV2, p < 0.0001; n = 9 per group. Data are mean ± SD. Panels B, D and F were compared across capsids by one-way ANOVA with Dunnett’s correction for multiple comparisons.

We next asked whether that gain is established at the cell surface. Vector was incubated with cells at 4 °C, a temperature that permits attachment but blocks endocytosis, and cell-associated genomes were quantified by ddPCR (Fig. 6C,D). On CHO-K1 the three capsids bound indistinguishably (p = 0.6217; p = 0.5238). On Lec8, AAV2.IvtB bound far more vector genomes per diploid genome than AAV2 (p < 0.0001), while AAV2.7m8 was unchanged (p = 0.3060). The transduction difference in Lec8 is thus already present at attachment.

Cell-based assays cannot separate a direct capsid–glycan interaction from an indirect consequence of altered glycosylation, so we tested binding to the glycan itself. Magnetic beads with or without immobilized GlcNAc were incubated with each vector and captured genomes were quantified as a percentage of input (Fig. 6E,F). AAV2.IvtB was recovered on GlcNAc beads roughly twice as efficiently as AAV2 (p < 0.0001), and AAV2.7m8 least of the three, identifying GlcNAc as a direct ligand of the AAV2.IvtB capsid.

Two further properties of the preparations bear on how these capsids behave *in vivo*. Differential scanning fluorimetry indicated that the VR-VIII insertions selectively lowered capsid-shell stability at neutral and early-endosomal pH (AAV2.IvtB Tm 62.9 °C at pH 7, vs 75.3 °C for AAV2) while the temperature at which the packaged genome became dye-accessible was indistinguishable across all five capsids (Fig. S6), locating the destabilization in the shell rather than in genome packaging. Particle composition was characterized in parallel: the fraction of genome-containing (full) particles ranged from 14.2% to 58% across capsids, lowest for AAV2 and highest for AAV2.IvtB (Fig. S7).

## Discussion

Intravitreal delivery would make AAV gene therapy for inherited retinal disease far safer and more broadly applicable than the subretinal injection on which current approaches depend, but the inner retina has remained a stubborn barrier to reaching photoreceptors from the vitreous.^2,29^ Most engineered capsids that improve intravitreal transduction have been selected for performance in a single species, usually mouse, and their advantage often attenuates in larger eyes.^7,30^ Here we took the inverse approach: rather than selecting for an outcome in one host and hoping it would generalize, we first identified the molecular features a vector must navigate, asked whether those features are conserved, and then engineered against them. The resulting capsid, AAV2.IvtB, enhanced photoreceptor transduction in both mouse and pig and conferred therapeutic benefit in two mechanistically distinct disease models.

The conceptual core of this work is that translatability can be designed in at the level of the target rather than discovered after the fact. By mapping the retinal distribution of heparan sulfate and terminal GlcNAc across seven mammalian species, we found that the two features relevant to intravitreal delivery, an HS-rich inner limiting membrane and terminal GlcNAc enrichment on photoreceptor segments, are conserved from mouse to human and most pronounced in primates. A capsid built to bind HS weakly while engaging terminal GlcNAc strongly is therefore anchored to targets that exist in the same configuration across the species in which it will eventually work. This reframes cross-species translation from a property one tests for into a constraint one selects under, and it offers a principle that extends beyond the eye: where the molecular barriers and targets of a delivery route are conserved, capsid engineering can be directed at those invariants rather than at species-specific surrogates.

This principle motivated CReST, a two-stage selection performed entirely *in vitro*. Decoupling capsid discovery from animal screening addresses a practical bottleneck, but its more important consequence is mechanistic transparency. Because each stage selects for a defined molecular property, the trajectory of a variant through the selection reports on its binding mode. AAV2.IvtA, AAV2.IvtB and AAV2.IvtC followed distinct paths: AAV2.IvtB remained near baseline through the HS stages and rose only upon GlcNAc selection, identifying it as a variant whose advantage rests on GlcNAc engagement rather than on altered HS binding; consistent with this, it matched parental AAV2 for both transduction and binding on the heparan-sulfate-bearing CHO-K1 background (Fig. 6B,D). The capsid advanced here therefore realizes the GlcNAc half of the design principle, and we have not measured its heparan-sulfate affinity directly. Direct testing supported this assignment: AAV2.IvtB transduced and bound Lec8 cells, which present N-glycans truncated to terminal GlcNAc, more efficiently than AAV2, was recovered on immobilized GlcNAc in a cell-free pull-down. Because enrichment at each stage is tied to a defined molecular property, the mechanism underlying a variant’s selection can be framed as a testable hypothesis rather than inferred from its rank in the output.

Across the pig retina and both therapeutic models, the advantage of AAV2.IvtB was expressed predominantly as greater transgene output per vector genome rather than as greater genome deposition. In the murine eye the relationship was inverted: AAV2.IvtB achieved the highest retinal vector-genome load at a transcript-per-genome ratio comparable to AAV2.7m8 (Fig. 3B). This species-dependent dissociation is consistent with, although it was not predicted by, the barrier model that motivated the selection: where intravitreal diffusion distances are short, as in the mouse, escape from the inner retina should translate directly into genome deposition, whereas in the larger porcine eye, where arrival at the outer retina is more constrained, differences in per-genome productivity would be expected to dominate. In the pig retina, the rd10 model and the laser-induced CNV model, AAV2.IvtB matched or fell below the benchmark capsids in vector-genome abundance while yielding the highest transgene transcript levels, the greatest preservation of outer nuclear layer thickness in rd10 retinas and the largest reduction in CNV lesion number. This dissociation between genome deposition and productive expression implies that the engineered tropism improves not only where particles arrive but how efficiently the delivered genomes are converted to product, plausibly through more productive uptake and trafficking at the photoreceptor segments, where terminal GlcNAc is concentrated. It also carries a practical implication: a capsid that achieves more per genome could in principle deliver equivalent benefit at lower dose, an attractive property for an intravitreally administered vector, where dose, immunogenicity and cost are linked. Enhanced output per genome could also arise downstream of receptor engagement, during intracellular processing of the capsid. Consistent with this, differential scanning fluorimetry showed that at neutral pH the AAV2.IvtB capsid is less thermostable than AAV2 while the thermostability of its packaged genome is unchanged (Fig. S6), consistent with a more flexible protein shell that retains its genome. Whether reduced shell stability accelerates uncoating cannot be determined from these data, because the two fluorimetry readouts report different molecular events and neither resolves kinetics. A direct test would require a DNase-protection genome-ejection assay after thermal or low-pH challenge, time-resolved uncoating measurements, or cryo-EM under endosomal-mimetic conditions.

A further observation concerns the interaction of AAV2.IvtB with the humoral immune system. After intravitreal delivery in naive animals, anti-AAV2 NAb titres in pigs were lower after AAV2.IvtB than after AAV2 or AAV2.7m8 in both serum and vitreous, with the larger difference in the vitreous (Fig. 4E,F). With one animal per capsid for serum and three eyes per capsid for vitreous, this is a descriptive observation. One possible structural explanation is that the 12-residue insertion in the VR-VIII loop, the region that contributes the immunodominant neutralizing epitopes of AAV2,^31,32^ reshapes the antigenic surface. The assay measured neutralization of wild-type AAV2 rather than of AAV2.IvtB, so a genuinely smaller response cannot be separated from one redirected away from shared epitopes; it was also performed in animals without prior AAV exposure and therefore speaks to *the de novo* antibody response rather than to the pre-existing anti-AAV2 immunity prevalent in humans.^33^ Because the preparations differed in full-particle fraction, the comparison is also confounded by the lower total capsid-antigen load AAV2.IvtB delivered at a matched genome dose. Resolving these will require matched-total-particle dosing and homologous neutralization measurements. Retinal transcript levels of IFN-γ, IL-1β, IL-6 and TNF-α also did not differ detectably among capsids in the same pigs (Fig. S4), giving no indication of a greater local inflammatory response at the dose tested.

The two therapeutic models test complementary requirements. Partial preservation of the outer nuclear layer in rd10 mice shows that AAV2.IvtB can deliver a corrective gene to the photoreceptors that degenerate in inherited retinal disease, the application that most needs an intravitreal route to photoreceptors. The progressive, time-dependent suppression of CNV shows that the same capsid can sustain expression of a secreted therapeutic, with lesion divergence emerging only at later timepoints in a manner consistent with durable gene-based rather than transient drug delivery.

Several limitations bound these conclusions and define the next steps. The rd10 experiment used a single endpoint: structure was assessed only at P45, so we cannot determine whether degeneration is slowed or arrested, and additional time points will be needed to establish the trajectory. Nor did we assess whether the spared photoreceptors are functional, so electroretinography or behavioral readouts will be needed to establish functional rescue. The pig data, while confirming the direction and significance of the effect in a large eye, showed a smaller fold-improvement than in mice, as expected from the greater diffusion distances; the porcine advantage was also distributed across several inner-retinal cell types rather than confined to photoreceptors, and whether this reflects a true species difference in tropism or a lower porcine baseline warrants direct study. Our immunological assessment remains incomplete: the cytokine panel is narrow, neutralization was measured against wild-type AAV2 rather than against AAV2.IvtB, and the full-particle confound noted above means a comprehensive evaluation (matched total-particle dosing, homologous neutralization, and T-cell responses) has yet to be completed. Finally, although terminal GlcNAc localization and our mechanistic assays extend to human tissue and human-derived cells, efficacy in the human eye remains to be demonstrated, and the molecular identity of the photoreceptor-segment terminal GlcNAc species engaged by AAV2.IvtB has not been resolved.

Beyond the specific capsid, this work offers a transferable template: characterize the conserved molecular barriers and targets of a delivery route, select against and toward them in defined *in vitro* assays, and validate the resulting mechanism across species. Applied to other tissues whose access barriers rest on conserved glycans or receptors, a receptor-guided, animal-free strategy of this kind could shorten the path from capsid discovery to a candidate with a defensible expectation of translating to humans.

## Materials and Methods

### Study approval and animals

All animal procedures were approved by the Institutional Animal Care and Use Committee of UMass Chan Medical School and conducted in accordance with the ARVO Statement for the Use of Animals in Ophthalmic and Vision Research. Adult C57BL/6J mice (5–8 weeks; The Jackson Laboratory) were used for capsid evaluation and the CNV model, and homozygous rd10 mice (B6.CXB1-Pde6b^rd10^/J; The Jackson Laboratory, stock 004297) for the photoreceptor-rescue study. Mice of both sexes were used and distributed across groups; animals were maintained on a 12-h light/dark cycle at 21–23 °C and 35–46% relative humidity with irradiated chow and water ad libitum. Domestic pigs (6 weeks old at injection; Yorkshire breed, purchased from Earl Parsons) were used for large-eye validation. Nonhuman-primate and human retinal tissue sections were donated control sections from a different study that was conducted under institutionally approved protocols ^34^; human tissue was de-identified post-mortem material from Department of Ophthalmology & Visual Sciences at UMass Chan, with post-mortem interval recorded. Animals were assigned to capsid groups at random, and all histological assessment were performed by observers masked to group identity. Predefined exclusion criteria (lens injury, vitreous hemorrhage, endophthalmitis or failed injection) were applied before analysis; the number of eyes and animals analyzed per group is stated in each figure legend.

### Retinal tissue from multiple mammalian species

To test whether HS and terminal GlcNAc distributions are conserved, retinas were collected from seven mammalian species: mouse, rat, rabbit, dog, pig, rhesus macaque and human. Globes were enucleated, fixed in 4% paraformaldehyde, cryoprotected in 30% sucrose, embedded in OCT and cryosectioned at 12 µm through the central retina; primate and human sections were taken from the macular region where available. Sections from all species within a given glycan comparison were processed in a single batch to permit cross-species intensity comparison.

### Immunohistochemistry

Retinal cryosections were air-dried, rehydrated in PBS (5 min), and ringed with a hydrophobic barrier pen. Sections were permeabilized in PBS containing 0.3% Triton X-100 (PBST) for 1 h and blocked in PBST with 5% BSA for 1 h at room temperature. Primary antibodies and lectins were diluted in blocking buffer and applied overnight at 4 °C. For cross-species glycan comparisons, sections were stained for heparan sulfate (anti-HS, clone F58-10E4, 1:100; AMSBIO) and terminal N-acetylglucosamine (succinylated wheat-germ agglutinin sWGA; FL-1021S, 1:100; Vector Laboratories); succinylation restricts WGA reactivity to terminal GlcNAc and minimizes binding to sialic acid. For transduction mapping in GFP-injected eyes, sections were stained for GFP (chicken anti-GFP, ab13970, 1:2,000; Abcam) in combination with a single cell-type marker per section: directly conjugated peanut agglutinin (PNA from Arachis hypogaea; L32460, 1:500; Thermo Fisher) for cone photoreceptor segments, mouse anti–glutamine synthetase (GS; MAB302, 1:1,000; Merck Millipore) for Müller glia, rabbit anti-BRN3A (ab245230, 1:200; Abcam) for retinal ganglion cells, or mouse anti-Chx10 (sc-365519, 1:200; Santa Cruz Biotechnology) for bipolar cells. Sections were washed three times in PBST (5 min each) and incubated for 2 h at room temperature with species- and isotype-matched Alexa Fluor–conjugated secondary antibodies (1:1,000; anti-chicken, anti-rabbit and anti-mouse IgG secondaries); directly conjugated PNA required no secondary. Nuclei were counterstained with DAPI (1:2,000; D9542, Sigma-Aldrich). After three further PBST washes (5 min each), sections were mounted and stored at 4 °C until imaging. Images were acquired on a Leica DM6 Thunder microscope fitted with a 16-bit monochrome camera using LAS X software, with acquisition settings held constant across all sections compared within an experiment.

### Quantification of retinal cell transduction

Quantification of photoreceptor transduction in retinas treated with vectors encoding cytosolic GFP under the CB6 promoter was performed by manually counting GFP-positive photoreceptor segments. For each group of injected mice, four rectangular regions of equal area surrounding the optic nerve head were selected from a single retinal section per eye. GFP-positive segments within these selected areas were manually counted. PNA was used to label cone outer segments. In addition, the total number of photoreceptor nuclei in the outer nuclear layer within the same areas was determined using the Imaris software package. Photoreceptor transduction efficiency was then calculated as the average per eye and per group by determining the ratio of GFP-positive photoreceptors to the total number of photoreceptors. Transduction of the remaining retinal cell types was quantified on the same four rectangular regions surrounding the optic nerve head, with each cell type scored on a separate section stained for its marker: glutamine synthetase for Müller glia, Brn3a for retinal ganglion cells and CHX10 for bipolar cells. Within each region, a masked observer manually counted the number of marker-positive cells co-expressing GFP and the total number of marker-positive cells, and transduction efficiency for each cell type was calculated as the average per eye and per group from the ratio of the two.

### AAV2 peptide-display library

A peptide-display library was constructed on the AAV2 *cap2* backbone by inserting a degenerate NNK cassette encoding a random 12-mer between residues N587 and R588 in the VR-VIII surface loop, a site that tolerates insertion without disrupting assembly or packaging. For the first round, the double-stranded NNK cassette, flanked by sequence overlapping the insertion site, was synthesized by Azenta and introduced by Gibson assembly into a *cap2* backbone plasmid, pAAV-Cap2-sc1, carrying AAV2 ITRs and a partial *Rep2* C-terminal sequence; full-length *Rep2* was supplied in trans during packaging from a Rep-only helper plasmid, pRep2-3stop, in which *cap2* is inactivated by three stop codons ^35^. The backbone was linearized by digestion with AflII and purified using a DNA and gel purification kit (D4008, Zymo Research); the linearized backbone and insert (backbone/insert = 1/5 ratio) were combined in a Gibson Assembly Master Mix reaction, and the resulting plasmid library was precipitated with isopropanol and concentrated in water. The library (<100 ng/µL) was electroporated into ElectroMAX DH10B cells (18290015, Invitrogen) following the manufacturer’s instructions; electroporated cells were pooled and cultured overnight in 500 mL of TB medium, and the plasmid library was then extracted and purified. For the second round of selection, VR-VIII sequences recovered from the first round were cloned by Gibson assembly into a fresh copy of the pAAV-Cap2-sc2 backbone, following the same linearization and assembly workflow, to generate the second plasmid library.

AAV library production was conducted in HEK293 cells ^35^. Calcium phosphate transfection was performed using 1.5 mg of pAd-DeltaF6 (containing adenovirus helper genes), 1.0 mg of pRep2-3stop, and 30 μg of the first or 100 μg of the second plasmid library in 10 roller bottles. Cells and culture medium were harvested 72 h post-transfection and pelleted by low-speed centrifugation; cells were lysed by three freeze-thaw cycles, and the supernatant was precipitated on ice with 40% polyethylene glycol solution (1:0.3 vol) followed by centrifugation. The lysate and supernatant were pooled and fractionated through four rounds of iodixanol gradient purification, with buffer exchange into PBS on Amicon-100 columns (Millipore). Titer was determined by ddPCR (QX200, Bio-Rad) against a poly(A)-specific primer/probe set, and capsid purity was assessed by silver staining of PAGE gels. Library quality at the plasmid and viral stages was assessed by amplicon deep sequencing of the VR-VIII variable region before selection (below). In the plasmid library, 85% of 91,360 filtered reads carried a full-length, in-frame 12-mer insert, per-position nucleotide frequencies matched the expected NNK distribution, and no single peptide exceeded 0.01% of reads, indicating uniform randomization without clonal dominance. Because an NNK 12-mer cassette spans a theoretical amino-acid space of approximately 4 × 10¹□, any sequencing depth undersamples the library; observed richness is therefore reported together with its sampling coverage rather than as a measurement of complexity.

### Matrigel/heparan sulfate barrier transduction assay

HeLa cells were seeded at 5 × 10^4^ cells per well in 24-well plates one day before infection. Transwell culture inserts (6.5 mm, 0.4 µm-pore polyester membrane; CLS3470-48EA, Corning) were coated with a Matrigel/heparan sulfate solution: Matrigel (CLS356231, Corning) was diluted in water to deliver 5 µg/cm², and heparan sulfate (HS; S5992, Selleckchem) was added at 0, 1, 50, 100, 150, 200, 250 or 300 µg per insert in a final coating volume of 70 µL per well. Coated inserts were incubated at 37 °C for 5 h, residual solution was removed, and the inserts were mounted into the wells over the seeded HeLa monolayer. A no-Matrigel, no-HS control received an uncoated insert. AAV2.GFP, AAV2.7m8.GFP or AAV9.GFP was then applied to the upper chamber at an MOI of 1 × 10^5^ vg per cell, and vector was left on the cells for the duration of the experiment. At 48 h post infection, GFP expression in the HeLa monolayer beneath the insert was imaged and quantified as mean GFP intensity per well. For transgene expression, total RNA was extracted from the monolayer at 48 h (DNA/RNA Purification kit; 47700, Norgen Biotek), reverse transcribed (iScript™ Reverse Transcription Supermix;1708841, BioRad), and GFP cDNA quantified by ddPCR and normalized to GUSB cDNA.

### CReST *in vitro* selection and sequencing

Selection was performed entirely *in vitro* in two stages. In the negative (HS-reduction) stage, HeLa cells were seeded at 3 × 10 cells per well in 6-well plates one day before selection. Transwell culture inserts (24-mm, 0.4-μm-pore polyester membrane; Corning) were coated with a Matrigel/heparan-sulfate solution: Matrigel (CLS356231, Corning) was diluted in water to deliver 5 μg/cm² (23.35 μg Matrigel per insert), and heparan sulfate (S5992, Selleckchem) was added to give 1,200 μg HS per insert, in a final coating volume of 280 μL per well. Coated inserts were incubated at 37 °C for 5 h; the residual solution was then removed, and the inserts were mounted into the wells over the seeded HeLa monolayer. The viral library (3 × 10^10^ vg) was applied to the upper chamber in 500 μL of culture medium together with adenovirus type 5 (Ad5) helper (MOI of 100). Only capsids that crossed the HS barrier reached and transduced the cells below; after 2 d, transducing variants were recovered by extracting HeLa total RNA (DNA/RNA Purification kit; 47700, Norgen Biotek), purifying capsid-derived mRNA (Dynabeads mRNA Purification Kit; 61006, Invitrogen) and reverse-transcribing with a cap2-specific primer (SuperScript IV First-Strand Synthesis System; 18-091-050, Invitrogen). To amplify the Cap sequence with 12-mer insertions, a nested PCR approach was used. In the first round of PCR, the target sequence was amplified using the initial set of primers (Nes-1F and Nes-1R) in multiple reactions containing 2 μL of cDNA, 25 μL of Q5 HotStart High-Fidelity 2× Master Mix (NEB), and a total reaction volume of 50 μL for 10 cycles. In the second round, amplification was performed with the second set of primers (Nes-2F and Nes-2R) in multiple reactions using 1 μL of amplicon purified and concentrated from the first-round PCR, 12.5 μL of Q5 2× Master Mix, and a total reaction volume of 25 μL for 20 cycles. The resulting amplicons were purified to generate second round of plasmid library before second round of AAV library production.

In the positive (GlcNAc-enrichment) stage, biotinylated N-acetylglucosamine (GlcNAc-biotin; G0297, TCI Chemicals) was dissolved in water to a 100 mM stock and diluted 10-fold immediately before use to 10 mM. The HS-reduced viral library (1 × 10^11^ vg) was mixed with 20 μL of 10 mM GlcNAc-biotin and incubated for 3 h at room temperature with shaking (1,200 rpm), in four replicate reactions per library to increase the final genomic DNA yield. Biotinylated GlcNAc–capsid complexes were then captured on Dynabeads MyOne Streptavidin C1 (65001, Invitrogen), 445 μL of beads per reaction, calculated from the manufacturer’s binding capacity of 20 μg biotinylated protein per 100 μL of beads and the 89 μg of GlcNAc present in each reaction. Beads were washed three times in Washing Buffer on a magnetic rack to remove unbound particles, and bound capsid genomes were recovered by genome extraction directly from the washed beads. The variable region was amplified as a 256 bp amplicon and sequenced at the MGH CCIB DNA Core using its CRISPR Sequencing service, an indexed amplicon deep-sequencing pipeline providing complete coverage of 200–280 bp amplicons. Observed richness was the number of distinct 12-mer peptide sequences. Reads were quality-filtered, translated and tallied, and a per-variant enrichment score was computed as the change in normalized frequency relative to the input AAV library. Variant abundance was tracked across the plasmid, HS-reduced library 1 (HS+) and GlcNAc-enriched library 2 (GlcNAc+) pools. AAV2.IvtA was advanced from the HS stage and AAV2.IvtB and AAV2.IvtC from the GlcNAc stage.

### rAAV production and titration

Single-stranded rAAV vectors encoding GFP, Pde6b or KH902 under a CMV-enhancer/chicken β-actin (CB6) promoter were produced in HEK293 cells by triple transfection with the ITR-flanked transgene plasmid, a Rep2/Cap plasmid encoding the relevant capsid (AAV2, AAV2.7m8, AAV2.IvtA, AAV2.IvtB, or AAV2.IvtC) and the adenoviral helper pAd-DeltaF6, using the same workflow as for the library. Cells were harvested 72 h after transfection, lysed, treated with benzonase, PEG-precipitated, purified over iodixanol step gradients and buffer-exchanged into PBS on Amicon-100 columns. rAAV titers were determined by droplet-digital PCR (ddPCR; QX200, Bio-Rad) with a transgene-specific primer/probe set, and capsid purity was confirmed by silver-stained SDS-PAGE showing the expected VP1:VP2:VP3 ratio. Vectors compared within an experiment were produced and titered in parallel to minimize batch effects. The proportion of genome-containing (full) versus empty particles was determined by negative-stain transmission electron microscopy particle counting (Fig. S7).

### Intravitreal injection in mice

Mice were anesthetized by intraperitoneal ketamine/xylazine (100/10 mg/kg) and pupils were dilated with topical phenylephrine and tropicamide. Using a pulled glass micropipette (Clunbury Scientific) introduced through the corneal limbus posterior to the ora serrata and angled to avoid the lens, 1 µL of vector containing 0.1% Fast Green was delivered into the vitreous under direct visualization. For capsid evaluation, mice received 2×10 vg/eye and were analyzed 4 weeks later. Eyes with lens injury, vitreous hemorrhage or reflux were excluded. For the neutralizing-antibody study, each animal received one capsid per eye.

### Intravitreal injection in pigs

Injections in pigs were performed as previously described ^36^. In brief, pigs were sedated and anesthetized, the ocular surface was disinfected with dilute povidone-iodine, and 80 µL of vector at 2×10^11^ vg/eye was injected into the mid-vitreous through the pars plana with an insulin needle, eyes were rinsed with sterile saline and collected 4 weeks after injection for analysis.

### Fundus and *ex vivo* imaging

Live GFP fundus fluorescence in mice was recorded under anesthesia on a Micron IV system (Phoenix Technology Group) at 4 weeks, with exposure and gain held constant across capsids. For pigs, the neural retina was dissected and imaged as posterior eyecups by fluorescence stereomicroscopy under matched acquisition settings. All comparative images within a figure were acquired and displayed with identical settings.

### Vector-genome and transcript quantification

Snap-frozen retinas were processed for parallel DNA and RNA recovery (DNA/RNA Purification kit; 47700, Norgen Biotek), and RNA was reverse-transcribed (High-Capacity cDNA RT Kit, Thermo Fisher). Vector genomes and transgene transcripts were measured by ddPCR against the transgene, GFP (Mr00660654_cn, Applied Biosystems), and a host reference gene (Tfrc (4458366, Applied Biosystems) for genome copies and Gusb (Mm01197698_m1) for transcripts in mouse; B2m for genome copies (Ss03391154_m1) and Gusb (Ss03387751_u1) for transcripts in pig). Genome copies were expressed per diploid genome; transgene transcripts were normalized to the host reference gene and are reported as fold change relative to the AAV2.7m8 benchmark. From porcine retinal RNA, IFN-γ (Ss03391054_m1), IL-1β (Ss03393804_m1), IL-6 (Ss07308316_g1) and TNF-α (Ss03391318_g1) transcripts were quantified by ddPCR, normalized to Gusb. A custom probe targeting the KH902 transgene was synthesized by Integrated DNA Technologies (IDT, Coralville, IA, USA) with the sequence 5′-TGAACTACGCCTGAGGATCCGATCT-3′.

### Cell lines

HeLa (ATCC CCL-2), used as the monolayer beneath the HS-coated Transwell in the negative-selection stage of CReST, were cultured in DMEM with 10% FBS. CHO-K1 (ATCC CCL-61) were cultured in F-12K medium (Kaighn’s modification of Ham’s F-12) with 10% FBS. The galactosylation-deficient mutant Lec8 (ATCC CRL-1737), a proline-auxotrophic Pro 5 derivative that lacks the Golgi UDP-galactose transporter and therefore displays N-glycans truncated to terminal N-acetylglucosamine (GlcNAc), was cultured in α-MEM with 10% FBS. All lines were maintained at 37 °C under 5% CO□.

### Transduction and flow cytometry

Cells were seeded in 48-well plates at 1.5 × 10^4^ cells per well and left to attach overnight. For all assays, vectors were diluted from a common master mix, applied in triplicate, and plates were incubated at 37 °C under 5% CO□. To define the glycan class required for attachment and transduction, CHO-K1 and Lec8 were transduced with GFP-expressing AAV2, AAV2.7m8, or AAV2.IvtB at a MOI of 1 × 10^5^, with adenovirus 5 as helper at MOI of 100; GFP expression was quantified at 48 h by fluorescence imaging and, in parallel cultures, by flow cytometry. Transduction efficiency was quantified in CHO-K1 and Lec8 cultures transduced in parallel with those used for imaging. At 48 h, the medium was removed, monolayers were washed once with PBS, dissociated with trypsin-EDTA, neutralized in complete medium and collected by centrifugation. Pellets were resuspended in PBS containing 2% FBS and filtered through a 40 µm cell strainer to obtain single-cell suspensions. GFP fluorescence was recorded in the FITC channel on a MACSQuant® Analyzer 16 (Miltenyi Biotec), acquiring 5,000 events per sample. Debris was excluded on forward- and side-scatter and doublets on pulse geometry, and the GFP-positive gate was set on untransduced cells of the same line processed in parallel. Mean GFP fluorescence intensity is reported.

### Binding assays

Vector attachment was examined in two complementary formats, a cell-based assay measuring surface binding and a cell-free pull-down measuring direct capsid–glycan interaction, both read out as vector genomes by ddPCR. Cell-surface binding was measured at 4 °C, which permits surface binding but prevents endocytosis, thereby separating attachment from post-entry steps. CHO-K1 and Lec8 were seeded in 24-well plates at 2.5 × 10^4^ cells per well and cultured overnight. Cells and vector stocks were pre-chilled at 4 °C for 1 h, GFP-expressing AAV2, AAV2.7m8 and AAV2.IvtB were added at MOI of 5 × 10^4^, and binding proceeded for 1 h at 4 °C. Unbound vector was removed by three washes with ice-cold PBS and cells were harvested. Total genomic DNA was extracted by phenol–chloroform–isoamyl alcohol (PCIA) precipitation, analyzed by ddPCR, with bound vector genomes quantified using a GFP-transgene probe and host genome copy number determined in parallel with a TFRC copy-number reference assay. Binding was expressed as vector genomes per diploid genome.

Direct binding of the capsids to GlcNAc was assessed in a cell-free streptavidin bead pull-down. A 100 mM stock of biotinylated GlcNAc (GlcNAc-biotin; G0297, TCI Chemicals) was prepared, aliquoted, and diluted to 10 mM before use. For each reaction, 1 × 10^11^ vg of AAV2.GFP, AAV2.7m8.GFP or AAV2.IvtB.GFP (20 µl) was combined with GlcNAc-biotin (20 µl; 89 µg) in a total of 40 µl (5 mM final) and incubated for 3 h at room temperature with shaking (1,200 rpm). Capsid–GlcNAc-biotin complexes were captured with streptavidin-coated magnetic beads for 30 min at room temperature with gentle rotation, collected on a magnet and washed four to five times with 100 µl PBS containing 0.1% BSA. For each capsid, three conditions were processed in parallel: an input sample (1 × 10^11^ vg, not subjected to capture) defining the total genomes added; a nonspecific-binding control incubated with streptavidin beads lacking GlcNAc-biotin; and the GlcNAc-binding sample captured on GlcNAc-biotin-conjugated beads. After the final wash, bead-associated genomes were released and purified by viral DNA extraction kit (12280050, Invitrogen) and quantified by ddPCR using a probe against the bovine growth hormone (BGH) polyadenylation signal. Specific binding was calculated for each capsid as (genomes bound to GlcNAc beads − genomes bound in the no-GlcNAc control) divided by input genomes, and expressed as the fraction of input vector genomes specifically captured.

### Differential scanning fluorimetry

The DSF assay was adapted from a previously published protocol ^37^. Briefly, SYPRO Orange (5,000×; Thermo Fisher Scientific) was diluted 1:100 in DPBS (Corning) to prepare a 50× working solution, and 45 µL of each AAV vector sample was mixed with 5 µL of the 50× SYPRO Orange solution. Fluorescence was monitored on a ViiA 7 Real-Time PCR System (Thermo Fisher Scientific): samples were equilibrated at 25 °C for 2 min, then ramped from 25 °C to 99 °C in 0.4 °C increments with a 2-min hold at each temperature, and fluorescence was recorded at each step using the ROX channel without a passive reference. To assess the effect of pH on capsid thermal stability, 5 µL of AAV vector, 5 µL of 50× SYPRO Orange and 40 µL of 0.6 M acetate buffer adjusted to pH 7.0–4.0 were combined before thermal analysis. The melting temperature (Tm) was defined as the temperature at the maximum first derivative of the fluorescence curve (−dF/dT) between 25 °C and 99 °C. To assess vector genome release, SYPRO Orange was replaced with SYBR Gold (Thermo Fisher Scientific) under otherwise identical conditions.

### AAV neutralizing antibody assay

Serum neutralizing antibody titers were determined using a cell-based luciferase transduction-inhibition assay. Huh7 cells were seeded into 96-well plates at 5 × 10 cells/well in 100 µL of complete DMEM (10% FBS) and incubated overnight at 37 °C and 5% CO to reach approximately 1 × 10 cells/well (90–100% confluence). Sera and vitreous from pigs were heat-inactivated and serially diluted two-fold in serum-free DMEM (1<1:5 to 1:10240). Each serum dilution (60 µL) was mixed with an equal volume of AAV2.Fluc, AAV2.7m8.Fluc or AAV2.IvtB.Fluc vector (60 µL), adjusted to deliver 1 × 10 vector genomes per well (MOI = 1), and the serum–vector mixtures were incubated at 37 °C and 5% CO for 1 h to allow neutralization. After approximately 50 min, 60 µL of medium was removed from each well, and 100 µL of the corresponding serum–vector mixture was added to the cells. Following a further 1 h incubation at 37 °C and 5% CO□, 100 µL of DMEM containing 20% FBS was added to each well, and the plates were incubated overnight. Firefly luciferase activity was then measured according to the manufacturer’s instructions, and luminescence (relative light units, RLU) was recorded on a plate reader. For each sample, AAV incubated with negative control serum was defined as 100% transduction, and relative transduction at each dilution was calculated as the background-subtracted RLU of the sample divided by that of the negative serum control. The neutralizing antibody titer (NT50) was defined as the reciprocal of the highest serum dilution that inhibited AAV transduction by at least 50% relative to the negative control.

### rd10 rescue and ONL morphometry

rd10 mice received a single intravitreal injection of single-stranded AAV2.Pde6b, AAV2.7m8.Pde6b, AAV2.IvtB.Pde6b or AAV2.GFP at postnatal day 15 (5×10^9^ vg/eye), with age-matched wild-type retina as the structural reference. At postnatal day 45, eyes were collected for histology and molecular analysis. ONL thickness was measured by a masked observer on nuclear-counterstained cross-sections through the optic-nerve head, within the same four rectangular regions of equal area used for transduction quantification, selected from a single retinal section per eye; values from the four regions were averaged per eye before group comparison. Retinal Pde6b transcript and vector-genome copies were quantified by ddPCR as above.

### Laser-induced CNV and anti-VEGF delivery

Choroidal neovascularization was induced in C57BL/6J mice by laser photocoagulation at 6 weeks of age as previously described (3-5 burns per eye around the optic disc) to rupture Bruch’s membrane ^38^. Fundus fluorescein angiography (FFA) was performed 5 d post damage to establish the total number of successful leakage sites, since not all initial damage sites produced sufficient injury to Bruch’s membrane to generate neovascular pathology; the number of leakage sites recorded on day 5 for each vector served as the denominator for the percentage of remaining leakage sites at later time points. On day 5, mice were injected intravitreally with single-stranded AAV2.KH902, AAV2.7m8.KH902, AAV2.IvtB.KH902 or the AAV2.GFP control, which served as the benchmark for natural healing of damaged sites. FFA images were then acquired every 10 d until 35 d post damage (days 15, 25 and 35), and the remaining active leakage sites were counted by a masked observer. Retinal KH902 transcript and vector genomes were quantified at endpoint by ddPCR.

### Statistical analysis

Data are presented as mean ± SD unless otherwise stated. The test applied to each comparison, the reference capsid where one was designated, the correction for multiple comparisons and n are given in the corresponding figure legend. In outline, several capsids were compared against a single reference by one-way ANOVA with Dunnett’s correction, and against one another by one-way ANOVA with Tukey’s correction. Two-group comparisons, such as transduction with and without the heparan-sulfate barrier, used unpaired two-tailed Student’s t tests. Time-course and dose-series data were analyzed by two-way ANOVA with correction for multiple comparisons. Neutralizing-antibody titers, which are interval-censored at two-fold dilution steps, were summarized as geometric mean titers; given the small per-group sample sizes, these comparisons are descriptive and were not subjected to significance testing. In the porcine study, four of six animals received the same capsid in both eyes, so eyes within an animal are not fully independent; porcine comparisons should be read with that correlation in mind. An adjusted p < 0.05 was considered significant. Analyses were performed in GraphPad Prism v10.

## Acknowledgments

The authors thank all members of the Gao laboratory and the Viral Vector Core at UMass Chan.

## Author contributions

J.H.W. conceived and designed the study. J.H.W., M.C. and W.Z. performed experiments, analyzed data and wrote the manuscript. D.G., C.P., Y.Y., R.H. and Q.S. performed experiments and analyzed data. J.X., J.M.S., C.P., Q.X. and X.K. provided resources and reagents. J.H.W. and G.G. supervised the study, acquired funding and edited the manuscript (co-corresponding authors). All authors reviewed and approved the final manuscript.

## Data and materials availability

All data needed to evaluate the conclusions in the paper are present in the paper and/or the Supplementary Materials. Source data for all quantified figures are provided with this paper. The 12-mer insertion sequences of AAV2.IvtA, AAV2.IvtB and AAV2.IvtC are given in Fig. 2F and in patent application WO2026217642. All raw NGS datasets will be made available upon request. Plasmids and capsid-encoding vectors are available from the corresponding authors under a material transfer agreement.

## Supplementary Materials

**Fig. S1.**
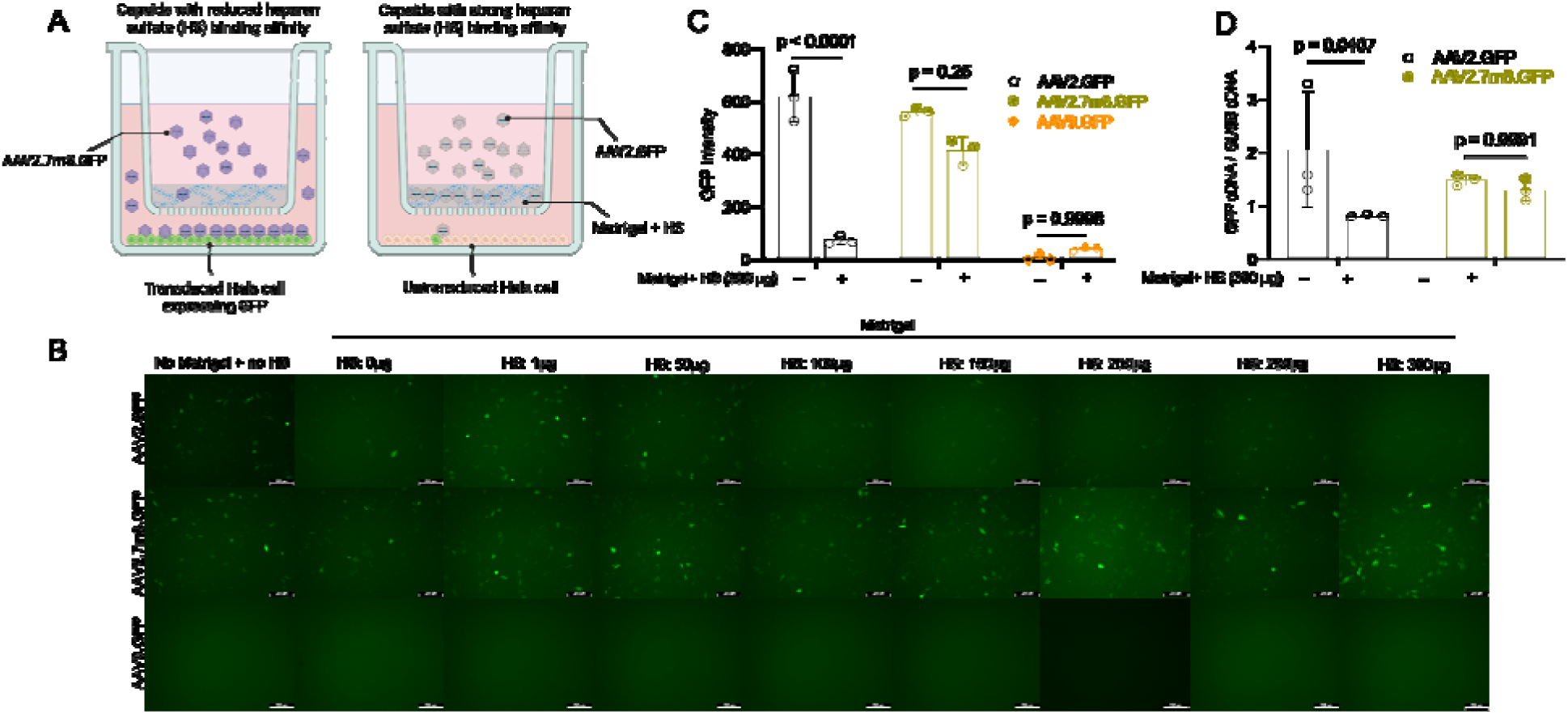
A heparan sulfate-loaded Matrigel barrier blocks AAV2 but not AAV2.7m8. **(A)** Assay design. HeLa cells are grown beneath a Matrigel layer supplemented with soluble heparan sulfate (HS). Capsids with strong HS-binding affinity (AAV2.GFP) are retained in the barrier and leave the cells untransduced, whereas capsids with reduced HS affinity (AAV2.7m8.GFP) cross it and drive GFP expression. (**B)** GFP fluorescence in HeLa cells 48 h after infection at an MOI of 1 × 10^5^ with AAV2.GFP, AAV2.7m8.GFP or AAV9.GFP, applied with no barrier or through Matrigel containing 0, 1, 50, 100, 150, 200, 250 or 300 µg HS. AAV2 signal falls away as HS is added, AAV2.7m8 persists at every HS load, and AAV9 stays at background throughout. (**C)** GFP intensity without (−) and with (+) Matrigel + HS (300 µg). The barrier almost abolishes AAV2 transduction (p < 0.0001), does not significantly reduce AAV2.7m8 (p = 0.25) and leaves AAV9, which transduces HeLa poorly, unchanged (p = 0.9998) (n = 3 per group). (**D)** GFP cDNA normalized to GUSB cDNA for AAV2.GFP and AAV2.7m8.GFP under the same conditions: the barrier lowers AAV2 expression (p = 0.0407) but not AAV2.7m8 (p = 0.9991) (n = 3 per group). Data are means ± SD. Statistical significance was calculated by unpaired two-tailed Student’s t test.

**Fig. S2.**
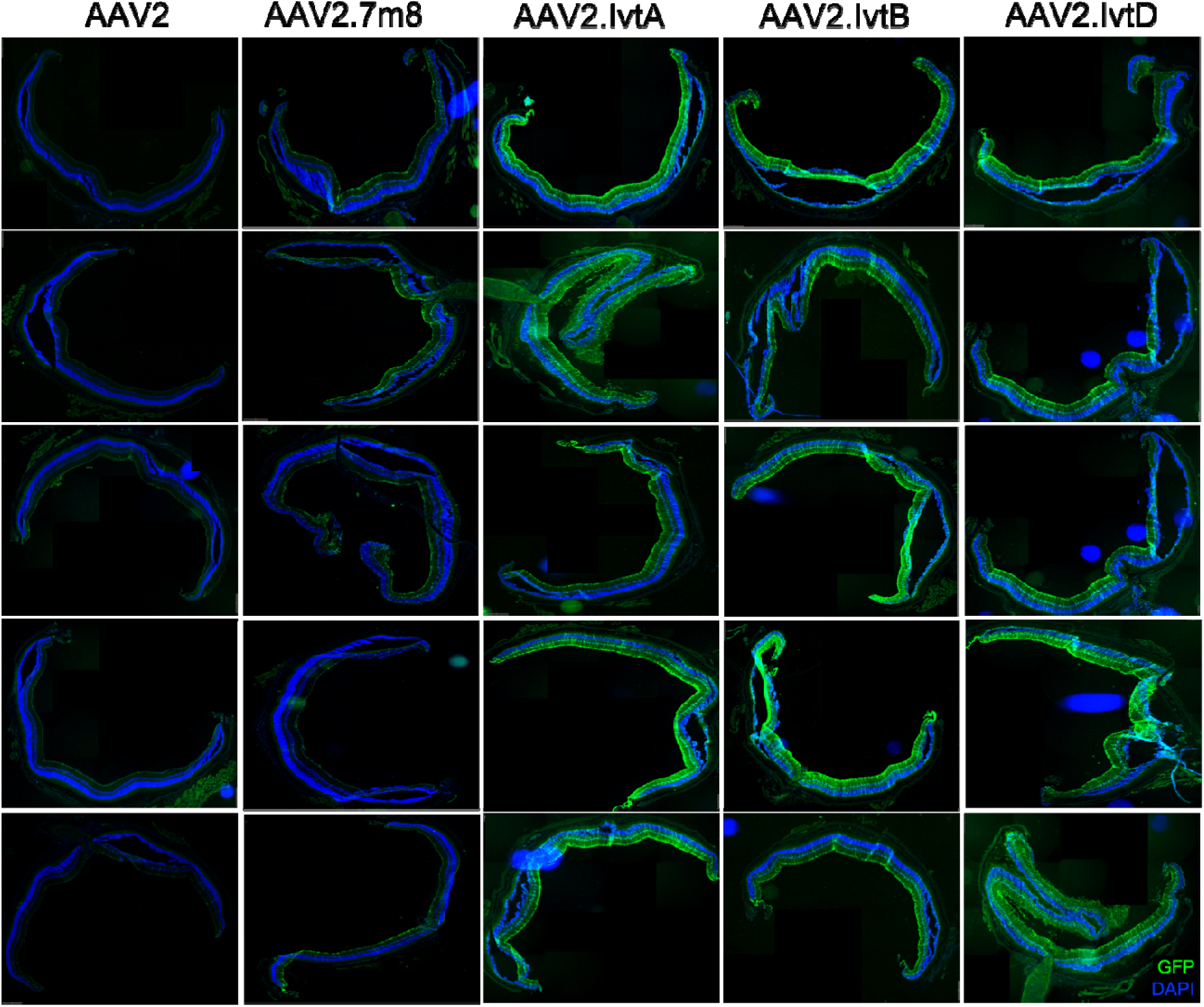
GFP expression across the mouse retina after delivery of AAV2 capsid variants. Retinal cryosections from mice injected intravitreally with a single-stranded CB6-GFP genome packaged in the AAV2, AAV2.7m8, AAV2.IvtA, AAV2.IvtB or AAV2.IvtC capsid (columns, left to right); each row is a separate eye (n = 5 per capsid). GFP, green; DAPI, blue. AAV2 eyes show only sparse GFP along a thin band of the inner retina; AAV2.7m8 shows a stronger GFP mostly in inner retinal cells; AAV2.IvtA, AAV2.IvtB and AAV2.IvtC give continuous GFP along the full length of the section in every eye, extending through the outer layers.

**Fig. S3.**
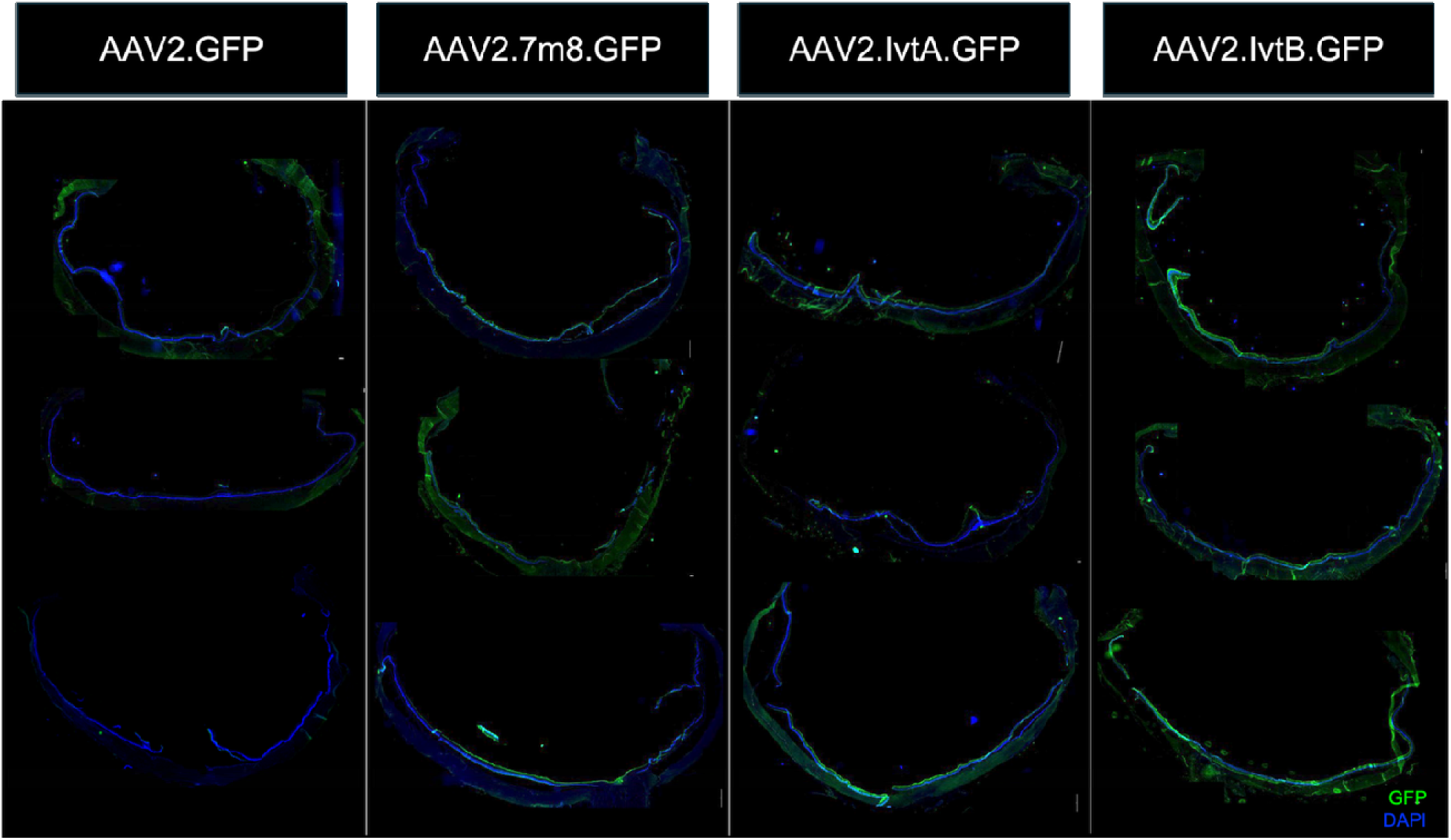
GFP expression in the pig retina after delivery of AAV2 capsid variants. Retinal cryosections from pigs injected with GFP vector packaged in the AAV2, AAV2.7m8, AAV2.IvtA or AAV2.IvtB capsid (columns, left to right); each row is a separate section (n = 3 per capsid). GFP, green; DAPI nuclear counterstain, blue. AAV2 sections show predominantly DAPI with only scattered GFP along the inner retina. AAV2.7m8 and AAV2.IvtA show GFP over part of the section, interrupted along its length. AAV2.IvtB gives the strongest and most continuous GFP, following the full length of th section. Domestic pigs (6 weeks old at injection) received 80 µL of vector at 2×10^11^ vg/eye into the mid-vitreous through the pars plana, and eyes were collected 4 weeks after injection. Sections were imaged on a Leica DM6 Thunder microscope with LAS X software, with acquisition settings held constant across all sections compared.

**Fig. S4.**
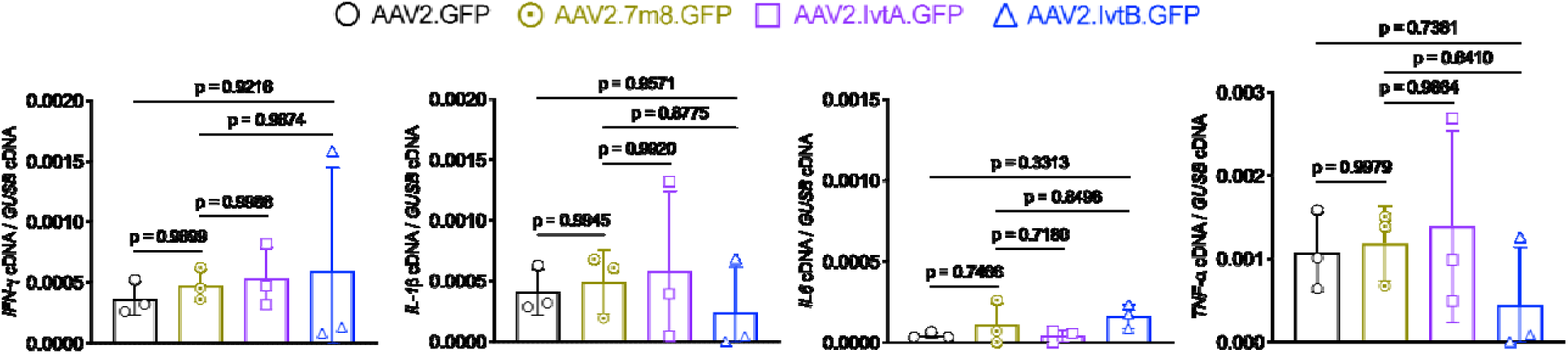
Intravitreal AAV2.IvtB does not produce a detectable retinal pro-inflammatory transcript signature in pigs. Pigs received a single-stranded CB6-GFP genome packaged in the AAV2, AAV2.7m8, AAV2.IvtA or AAV2.IvtB capsid, delivered intravitreally at 2×10^11^ vg/eye; these are the animals described in Fig. 4. Retinal transcript levels of IFN-γ, IL-1β, IL-6 and TNF-α were measured by ddPCR and normalized to Gusb (n = 3 per capsid). No significant difference was detected among capsids for any of the four transcripts. Data are mean ± SD; each transcript was compared across capsids by one-way ANOVA with Tukey’s correction for multiple comparisons.

**Fig. S5.**
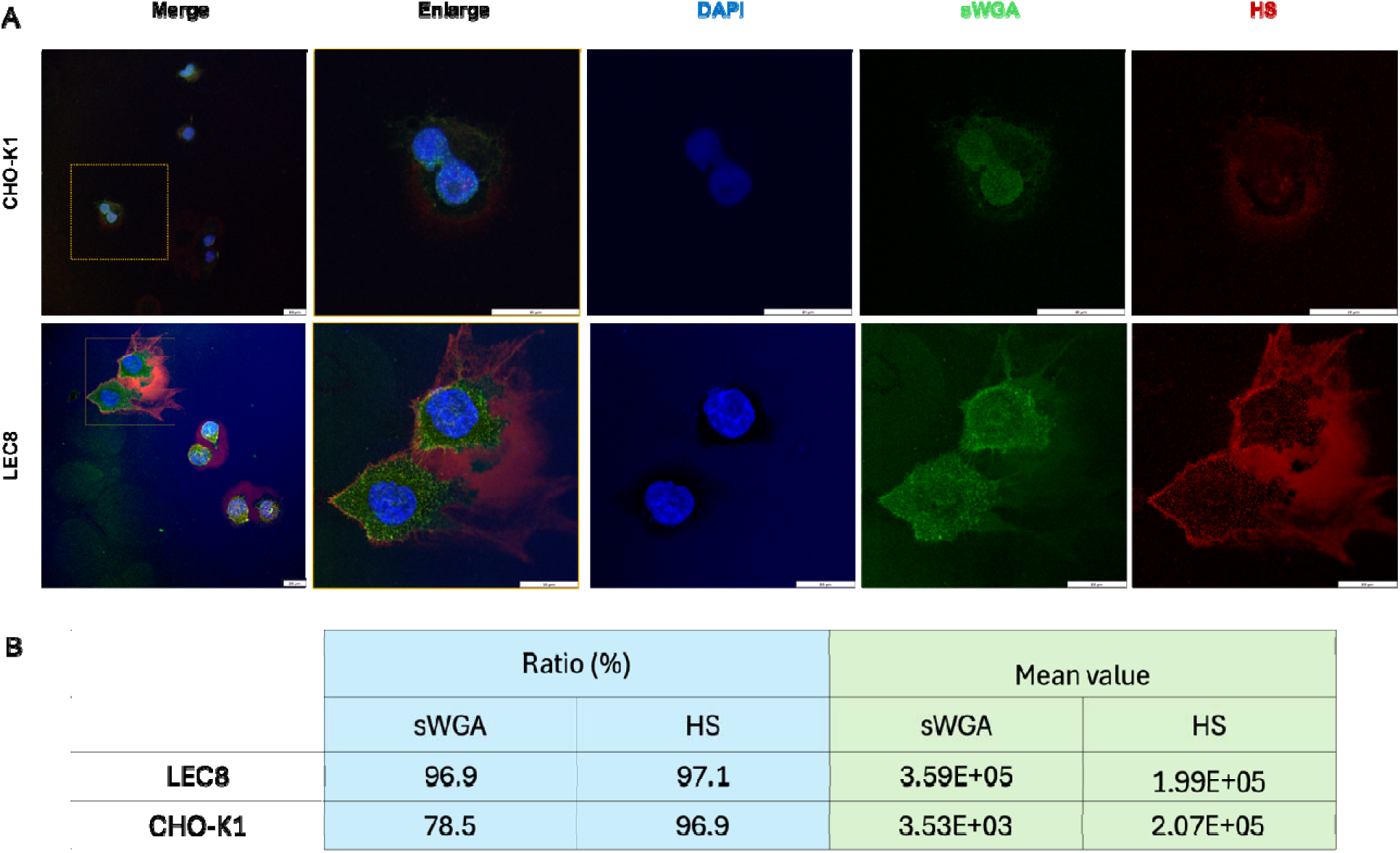
CHO-K1 and Lec8 cells differ in surface terminal GlcNAc but carry comparable heparan sulfate. (A) CHO-K1 (top row) and Lec8 (bottom row) stained for nuclei (DAPI, blue), terminal N-acetylglucosamine with succinylated wheat-germ agglutinin (sWGA, green) and heparan sulfate (HS, red). CHO-K1 shows weak, diffuse sWGA labeling, whereas Lec8 shows strong punctate sWGA at the cell surface, consistent with exposure of terminal GlcNAc on loss of the Golgi UDP-galactose transporter; HS labeling is present in both lines. Scale bars, 20 µm. (B) Flow cytometry of the same two lines stained for sWGA and HS. Ratio (%), percentage of positive cells; mean value, mean fluorescence intensity. Both lines were predominantly sWGA-positive (Lec8 96.9%, CHO-K1 78.5%), but sWGA mean fluorescence differed by two orders of magnitude (Lec8 3.59 × 10 versus CHO-K1 3.53 × 10³). HS was comparable between the lines in both positivity (Lec8 97.1%, CHO-K1 96.9%) and mean fluorescence (1.99 × 10 and 2.07 × 10□). The two lines therefore differ in exposed terminal GlcNAc on a shared heparan-sulfat background.

**Fig. S6.**
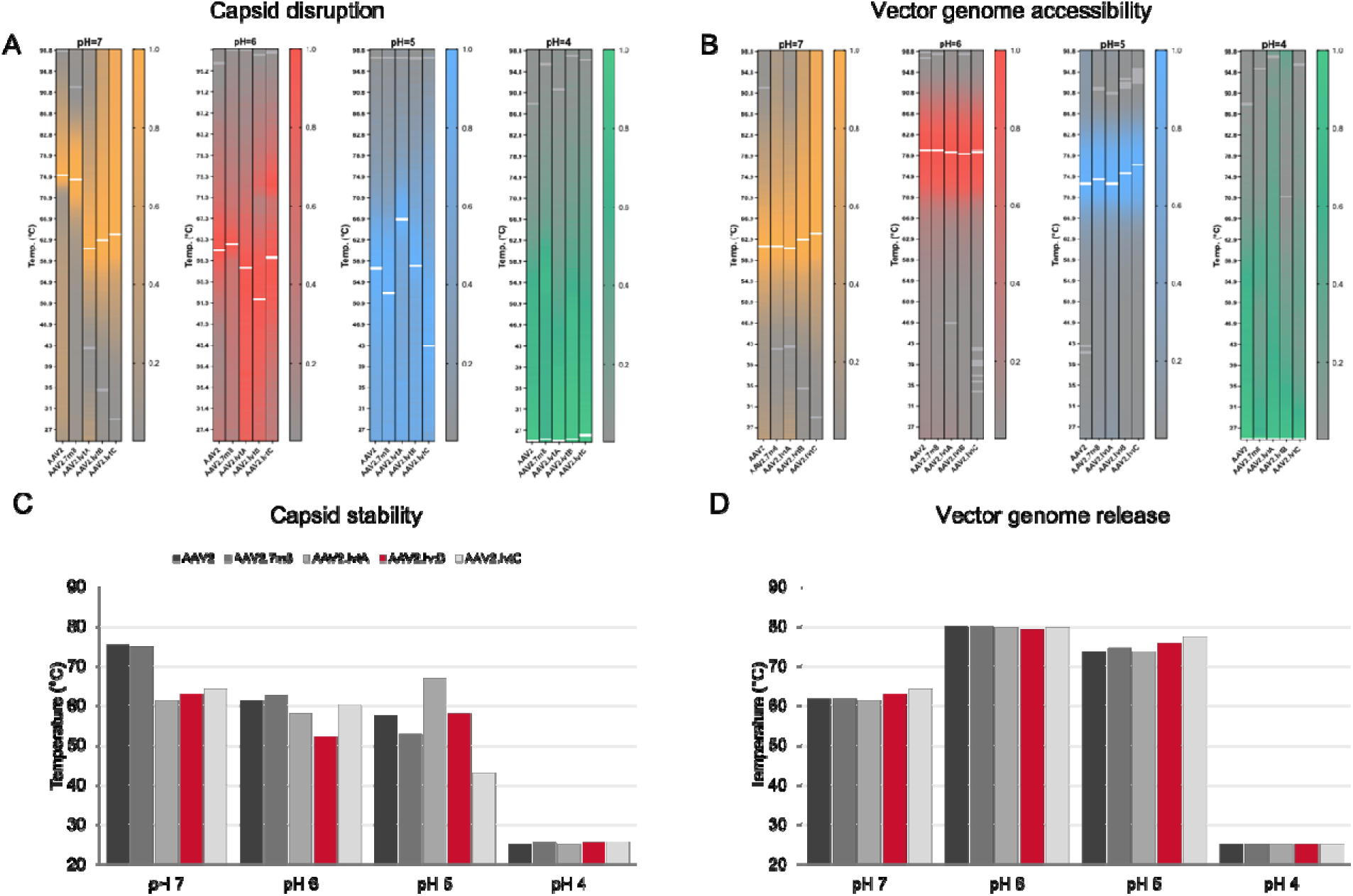
Differential scanning fluorimetry of AAV2 capsid variants across the endosomal pH range. Purified AAV2, AAV2.7m8, AAV2.IvtA, AAV2.IvtB and AAV2.IvtC were heated from 25 to 99 °C at pH 7.0, 6.0, 5.0 and 4.0, with SYPRO Orange reporting capsid protein unfolding and SYBR Gold reporting exposure of the packaged genome. (A and B) Heatmaps of capsid disruption (A) and vector genome accessibility (B) as a function of temperature; within each pH panel, lanes are AAV2, AAV2.7m8, AAV2.IvtA, AAV2.IvtB and AAV2.IvtC from left to right, and color intensity denotes signal relative to each lane’s peak (scale 0 to 1). (C and D) Melting temperatures (Tm, the temperature at the maximum first derivative of the fluorescence curve) for capsid disruption (C) and vector genome release (D). Capsid stability was both capsid- and pH-dependent: at pH 7 AAV2.IvtB disrupted at 62.9 °C, about 12 °C below AAV2 (75.3 °C) and AAV2.7m8 (74.9 °C); at pH 6 it was the least stable variant (52.1 °C versus 61.3 °C for AAV2); and at pH 5 it was comparable to AAV2 (58.1 versus 57.7 °C), where AAV2.IvtA was the most stable variant (66.9 °C) and AAV2.IvtC the least (42.9 °C). Vector genome release was similar across all five capsids at each pH (pH 7, 61.3 to 64.1 °C; pH 6, 79.3 to 80.1 °C; pH 5, 73.7 to 77.3 °C), so the reduced capsid stability of AAV2.IvtB at neutral and early-endosomal pH is not accompanied by earlier genome extrusion. At pH 4 both readouts reached their maximum at the lowest temperature measured (25 °C). Because SYPRO Orange and SYBR Gold signals are themselves pH-sensitive, this condition was not interpreted further.

**Fig. S7.**
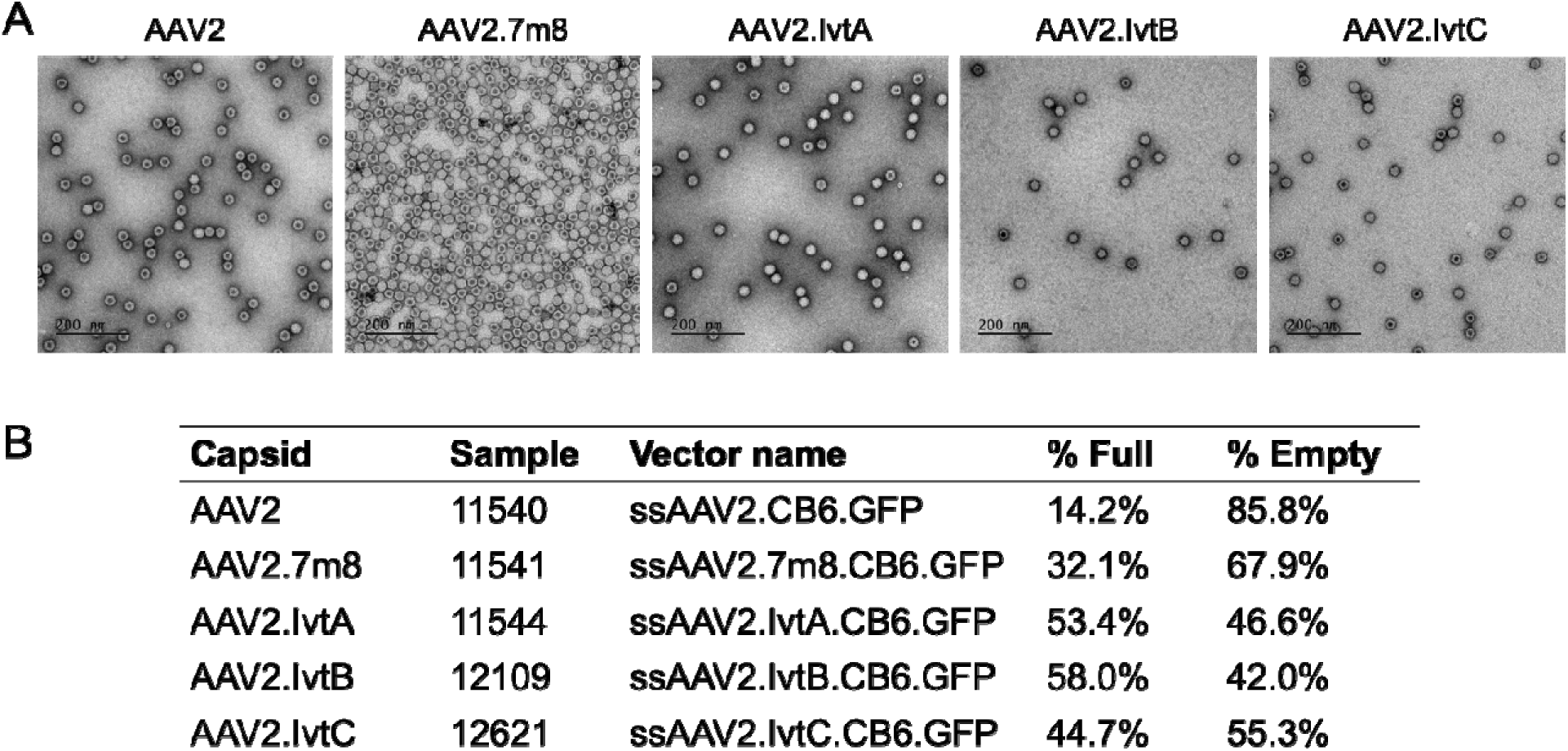
Particle morphology and genome packaging of the AAV2 capsid variants. (A) Negative-stain transmission electron micrographs of purified single-stranded (ss) AAV2, AAV2.7m8, AAV2.IvtA, AAV2.IvtB and AAV2.IvtC vector. Particles with a stain-filled centre are empty capsids; uniformly dark particles are genome-containing (full) capsids. All five preparations show intact, uniform particles of the expected size with no aggregation or debris. Scale bars, 200 nm. (B) Full and empty particle ratios for th same preparations, each packaging the CB6.GFP genome. Full-capsid content rises from 14.2% for AAV2 and 32.1% for AAV2.7m8 to 53.4% (IvtA), 58.0% (IvtB) and 44.7% (IvtC).

Supplementary Table S1. Heparan sulfate (HS□) affinity selection of the AAV2 N587 12-mer peptide-display library.

Supplementary Table S2. N-acetylglucosamine (GlcNAc□) affinity selection of the AAV2 N587 12-mer peptide-display library.

